# A membrane-impermeant nucleic acid dye converts bacteriophage plaque assays into a machine-readable format for automated counting

**DOI:** 10.64898/2026.08.07.741843

**Authors:** Alissa Wiwi, Jack Arnold, Darren W. Branch, Jesse Cahill

**Affiliations:** Biological & Chemical Sensors, Sandia National Laboratories, Albuquerque NM, USA; Environmental Systems Biology, Sandia National Laboratories, Albuquerque NM, USA

**Keywords:** bacteriophage, plaque assay, fluorescence, ImageJ, automated image analysis

## Abstract

Plaque assays remain the gold standard for bacteriophage quantification, but routine plaque counting is labor-intensive, time-consuming, and poorly suited to large experiments or automated workflows. Conventional plaque images also often provide insufficient contrast for simple software-based counting, especially when plaques are small, faint, or heterogeneous. Here we show that a membrane-impermeant nucleic acid dye (SYTOX) can convert standard bacteriophage plaque assays into a high-contrast, machine-readable format compatible with simple automated counting. In a soft-agar overlay workflow, fluorescent labeling enabled plaque detection and automated enumeration using an open-source ImageJ pipeline based on Find Maxima. In a representative benchmark using a large language model (LLM), conventional transmitted-light images were not counted accurately, whereas fluorescent SYTOX images brought both Fiji- and LLM-based counts within Poisson counting uncertainty of the manual count. Because the method improves the image contrast of the assay itself, it may also provide improved input for future machine-learning or other advanced automated counting workflows. The method was evaluated across diverse phage-host systems spanning dsDNA, ssRNA, filamentous, and enveloped phages, including T7, MS2, M13, and phi6. In lytic systems, fluorescent signal emerged prior to or alongside conventional plaque visibility and yielded automated counts that agreed closely with manual counting. M13 exhibited delayed fluorescence consistent with its chronic, nonlytic lifestyle, yet remained machine-countable at the conventional next-day endpoint. A Gram-positive Leo2–*Bacillus safensis* system revealed an important compatibility limit: dye incorporation at plating inhibited plaque formation. Follow-up experiments indicated that SYTOX did not measurably reduce free Leo2 recovery or alter host-dependent depletion of free phage in liquid culture, arguing against simple direct phage damage or a large liquid-phase adsorption defect under the tested conditions. While we were unable to identify the exact cause of the plaquing defect, a post-labeling workflow restored detectability and automated counting. Together, these results show that membrane-impermeant dye labeling can make plaque assays more computationally tractable while preserving the accessibility of standard phage methods. This approach provides a practical path toward higher-throughput, statistically rigorous phage biology in both low-resource and automation-oriented laboratories.

## 1 Introduction

In phage biology, plaque assays remain a foundational method to directly quantify infectious particles, support clonal isolation, and allow investigators to evaluate host range and infection phenotypes in a standardized way (Abedon and Yin 2009, Kropinski, Mazzocco et al. 2009, Acs, Gambino et al. 2020, Panteleev, Kulbachinskiy et al. 2025). Despite their central role, plaque assays retain a major practical limitation: counting plaques by eye is slow, repetitive, and difficult to scale. This becomes especially burdensome in experiments with multiple strains, phages, dilutions, and replicates. Moreover, the precision of plaque assays is fundamentally constrained by Poisson counting statistics, in which the uncertainty associated with a count is approximately ± the square root of N, where N is the number of plaques counted. Counts that fall within this interval are not statistically distinguishable and therefore represent equivalent estimates of phage titer. As a result, higher-precision experiments require larger numbers of plaques, more replicate plates, or both. These needs increase the manual burden of plaque counting and can quickly make otherwise straightforward experiments impractical at scale. In practice, manual counting can become the rate-limiting step that constrains experiment size, replication, and throughput (Kropinski, Mazzocco et al. 2009, Anderson, Rashid et al. 2011, Cacciabue, Curra et al. 2019).

This limitation is increasingly important because phage research is moving toward larger and more quantitative workflows. Recent reviews highlight growing interest in high-throughput plaque methods, robotic screening, and modern quantitative assays for phage-host interactions (Xie, Wahab et al. 2018, Storms, Teel et al. 2020, Haines, Hodges et al. 2021, Egido, Toner-Bartelds et al. 2023, Panteleev, Kulbachinskiy et al. 2025). However, standard plaque assays remain visually optimized for human readers rather than for software. Conventional white-light plaque images often contain low-contrast clearing against heterogeneous lawns, irregular edges, and morphology that depends strongly on growth conditions, agar composition, host physiology, and phage biology (Abedon and Yin 2009, Panteleev, Kulbachinskiy et al. 2025). Even where automated plaque-analysis tools exist, their performance can be limited by plaque contrast and morphology, particularly for faint or turbid plaques, and preprocessing is often still required (Yakimovich, Andriasyan et al. 2015, Katzelnick, Coello Escoto et al. 2018, Cacciabue, Curra et al. 2019, Trofimova and Jaschke 2021, Panteleev, Kulbachinskiy et al. 2025).

A conceptually attractive solution is to shift plaque detection from contrast-based readout to fluorescent signal-based readout. Membrane-impermeant nucleic acid dyes are a promising basis for such a strategy because they are largely excluded from intact cells but fluoresce strongly upon gaining access to nucleic acids after membrane compromise or lysis (Roth, Poot et al. 1997, Lebaron, Catala et al. 1998). In phage research, related approaches have already been used in liquid culture to monitor phage-induced lysis in real time (Egido, Toner-Bartelds et al. 2023), and more broadly, fluorescence-based viability and lysis assays have become useful tools for studying infection dynamics outside classical plaque formats (Low, Bohnlein et al. 2020, Panteleev, Kulbachinskiy et al. 2025). In eukaryote virology, fluorescent plaque assays using DNA dyes have also been shown to improve temporal resolution, enable automated plaque analysis, and support live-cell imaging of cytopathic effects (Yakimovich, Andriasyan et al. 2015, Culley, Towers et al. 2016, Katzelnick, Coello Escoto et al. 2018, Arias-Arias, Corrales-Aguilar et al. 2021). Together, these studies suggest that nucleic-acid-access dyes can report infection-associated cell damage well before conventional endpoint visualization.

However, to our knowledge, this strategy has not been established as a practical method for standard bacteriophage agar-overlay plaque assays specifically aimed at improving simple automated plaque counting across diverse phage-host systems. Here we sought to develop a practical fluorescent plaque-labeling strategy that would make bacteriophage plaque assays machine-readable without sacrificing the simplicity and accessibility that make plaque assays so widely used. In particular, we aimed for a method that could support simple automated counting in standard laboratories today while also providing a foundation for more highly automated phage workflows in the future. To be broadly useful, the method should rely on standard plaque assay materials, avoid phage engineering, require no custom imaging instrumentation or machine-learning pipeline, and improve software detectability enough to justify adoption. Using this framework, we tested whether fluorescent plaque labeling could provide an accessible route to higher-throughput, automation-ready bacteriophage quantification without phage engineering or custom imaging infrastructure

## 2 Methods

### 2.1 Bacterial strains, bacteriophages, and culture conditions

Bacteriophages T7, MS2, M13, phi6, and Leo2 and their corresponding bacterial hosts were propagated and handled using standard procedures as described previously by our lab (Humphrey, Mackenzie et al. 2023, Humphrey, Tezak et al. 2023). High-titer lysates were prepared in advance and stored at 4 °C until use. Host cultures were initiated from isolated colonies and grown overnight in LB broth under host-appropriate conditions. *Escherichia coli* hosts for T7, MS2, and M13 were cultured at 37 °C with aeration, *Pseudomonas syringae* for phi6 was cultured at 28 °C, and *Bacillus safensis* for Leo2 was cultured at 30 °C. For MS2 and M13 assays, overnight *E. coli* cultures were subcultured 1:100 v/v into fresh medium in 4 mL culture tubes and grown for approximately 2 h before use to provide actively growing cells for plaquing. Additional strain identity, media composition, and routine microbiological handling were described previously (Humphrey, Mackenzie et al. 2023, Humphrey, Tezak et al. 2023).

### 2.2 SYTOX plaque-labeling workflow

SYTOX Green membrane-impermeant nucleic acid dye Solution in DMSO, Thermo Fisher Scientific (Cat no. S7020), was used for fluorescent plaque labeling. Stock dye was maintained at 5 mM, aliquoted to minimize repeated freeze-thaw cycles, stored at -20 °C, and protected from light during storage and handling.

Plaque assays were performed using a standard soft-agar overlay workflow. Briefly, 100 µL of diluted phage lysate was mixed with 200 µL of the corresponding host culture and incubated for approximately 10 min to allow adsorption. The mixture was then added to 5 mL molten soft agar and poured onto the surface of a base agar plate. For fluorescent plaque assays, SYTOX was incorporated directly into the molten soft agar immediately before plating by addition of 5 µL of 5 mM stock per 5 mL overlay, yielding a final concentration of 5 µM. Matched no-dye controls were prepared in parallel without SYTOX addition. Plates were incubated under host-appropriate conditions as described above.

### 2.3 Post-labeling workflow for the Leo2–*Bacillus safensis* system

Because incorporation of SYTOX during plating inhibited plaque formation in the Leo2–*Bacillus safensis* system, an alternative post-labeling workflow was used. Leo2 plaque assays were first prepared under standard no-dye conditions. After plaque development, 25 µL of 5 mM SYTOX was diluted in 5 mL SM buffer to a final concentration of 25 µM and applied to the plate surface. Plates were agitated gently on a circular shaker for 30 min while covered, followed by 30 min uncovered, protected from light throughout. Plates were then allowed to dry before imaging.

### 2.4 Imaging

Plaque development was monitored by time-course imaging at the intervals indicated in the results section. Plates were imaged using a Syngene G:BOX Chemi XX6 imaging system controlled with GENESYS software. For fluorescence imaging, plates were imaged using the Alexa Fluor 500 setting with the Filt 525 emission filter and Blue LED Module (M) illumination. For transmitted-light imaging, plates were imaged using the Amido Black setting with the UV06 filter and upper white-light illumination. We also explored fluorescence imaging on a Gel Company UltraBright LED Transilluminator TLL-100 using 470 nm blue LED illumination and found that plaques remained machine-countable under this alternative setup (**Supplementary Figure 7**). Although not systematically benchmarked across all conditions, this result suggests that SYTOX-based plaque contrast should be transferable to other standard fluorescence imaging platforms commonly available in microbiology laboratories. After intermediate imaging, plates were returned to incubation for continued plaque development. Endpoint images were used to compare fluorescent signal emergence with conventional plaque visibility.

### 2.5 Manual and automated plaque counting

Plaques were counted manually by visual inspection (Schindelin, Arganda-Carreras et al. 2012). Manual counts were used as the reference for automated comparisons. Fluorescence images were analyzed in Fiji/ImageJ. Images were cropped to the plaque assay area, converted to 32-bit format, and analyzed using the Find Maxima function in Fiji/ImageJ. For transmitted-light images, image inversion was performed using the Light Background feature in Fiji/ImageJ. Prominence was initially explored over a practical range, typically 15-25, by visually assessing whether detected maxima localized to discrete fluorescent plaques rather than background variation. The guiding principle for this initial step was spatial plausibility of plaque-associated maxima rather than direct numerical agreement with the manual count. If lower prominence values produced excessive background detections or higher values failed to mark clearly visible plaque-associated puncta, prominence was adjusted to an intermediate value that best localized maxima to plaque signal while minimizing obvious false positives. Detected maxima were displayed as point selections, and automated plaque counts were recorded as the number of plaque-associated maxima.

### 2.6 Early fluorescent puncta assessment

To assess whether fluorescent puncta could be detected during early plaque development, plates were imaged by both fluorescence and transmitted light over time. Early fluorescent puncta, visible plaques, and their spatial correspondence at later time points were assessed by comparison of the image series. Endpoint images were used to determine whether early fluorescent puncta corresponded to mature plaque positions.

### 2.7 Plaque recovery

To test compatibility of fluorescent labeling with downstream phage recovery, well-isolated plaques were picked from SYTOX-labeled and matched no-dye control plates for T7, MS2, phi6, and Leo2. For T7, MS2, and phi6, plaques were recovered from plates in which SYTOX was present during plaque formation. For Leo2, plaques were recovered from plates labeled by the post-incubation workflow described above.

Individual plaques were aseptically cored using a transfer pipette, suspended in 1 mL SM buffer, and clarified by filtration through sterile 0.22 µm filters. Filtrates were serially diluted in SM buffer and re-titered by standard plaque assay using the corresponding host. Titers recovered from SYTOX-labeled plaques were compared with titers from matched no-dye controls to assess whether fluorescent labeling altered recovery of infectious phage.

### Liquid-phase Leo2–Bacillus safensis SYTOX interaction assay

To test whether SYTOX altered Leo2 behavior under liquid conditions prior to plaque formation, a short liquid-phase assay was performed using Leo2 with and without Bacillus safensis cells in the presence or absence of dye. Leo2 stock at 2.4 ×10^9^ PFU/mL was diluted 10^−5^ to a working concentration of 2.4 ×10^4^ PFU/mL. *B. safensis* was subcultured 1:50 into fresh LB and grown to an A_600_ of ∼0.4. 100 µL aliquot of the culture was removed for CFU determination, and the remaining culture was used immediately for the assay.

Host-containing reactions were prepared by combining 900 µL of B. safensis culture with 100 µL of the Leo2 working dilution, with or without 1 µL SYTOX using a 5 mM stock, resulting in a final concentration of 5 µM. Parallel no-cell controls were prepared in LB broth alone with the same phage input, with or without SYTOX. Reactions were mixed gently and held at room temperature. At 10 min and 30 min after mixing, 200 µL aliquots were removed and immediately centrifuged at maximum speed for 1 min. A 120 µL portion of the clarified top supernatant was carefully transferred to a fresh tube, serially diluted as needed, and plated by standard plaque assay to quantify supernatant-recoverable free phage.

### 2.1 Large language model-based plaque detection and quantification

Plaque counts were obtained via automated image analysis performed by Claude (Sonnet 5.5, Anthropic), using Python 3 with OpenCV and scikit-image. For each full-plate image (transmitted light or fluorescence), the plate boundary was detected by Gaussian smoothing, thresholding, and contour fitting, and a lawn region of interest (ROI) was defined at 90% of the detected plate radius to exclude specular reflection artifacts from the dish rim. Background illumination was flattened using morphological opening (41px structuring element) to estimate and subtract local background, and the resulting image was normalized within the lawn ROI.

Discrete plaques were identified as local intensity maxima using multi-scale Laplacian-of-Gaussian blob detection (σ range 3.5–12px, 8 scales). Detections within 18px of the ROI boundary were excluded to avoid residual rim artifacts (also applied to plate images used in Fiji/ImageJ comparison). Plaques with two resolvable intensity maxima were counted individually, even where boundaries overlapped; merged regions lacking a resolvable local minimum between centers were flagged as ambiguous. All automated detections were visually validated against source images prior to reporting.

The following prompt was used to perform this analysis in a reproducible manner: “Analyze the attached bacteriophage plaque assay image(s) (full Petri dish, soft-agar overlay; transmitted light or fluorescence). Detect the plate boundary and exclude the outer glass-rim reflection band and any region outside the lawn. Identify discrete plaques as localized clearings (transmitted light) or bright puncta (fluorescence), correcting for uneven background illumination. Count plaques with clearly separable centers individually, even if touching or overlapping; flag merged regions with indistinguishable centers as ambiguous. Exclude plaques more than half outside the image/lawn boundary and any non-plaque artifacts (dust, bubbles, scratches, labels, condensation). Report: total count, ambiguous count, confidence level, counting basis, brief image-quality notes, and approximate plaque coordinates (pixel or relative).”

### 2.2 Statistical analysis

Recovered phage titers from SYTOX-labeled and matched no-dye control plaques were compared separately for each phage using unpaired two-tailed t-tests in GraphPad Prism 10. Differences were considered statistically significant at p<0.05.

For the liquid-phase Leo2–Bacillus safensis assay, supernatant-recoverable phage titers were converted to log_10_ (PFU/mL) before analysis. Log-transformed values were analyzed by ordinary three-way ANOVA with factors of time, host-cell presence, and SYTOX presence, followed by Šídák’s multiple-comparisons test for planned pairwise comparisons. Differences were considered statistically significant at p<0.05.

## 3 Results

### 3.1 Fluorescent labeling converts plaque assays into a high-contrast machine-readable format

We first tested whether fluorescent labeling improved detectability of plaques relative to conventional plaque imaging. In the T7 system (Dunn and Studier 1983), plates prepared with dye showed clearly localized fluorescent signal within plaques, whereas matched dye-free plates imaged under the same fluorescence conditions showed no corresponding signal (Figure 1A). In contrast, standard visible-light images of matched dye-containing plates were not obviously distinguishable from dye-free plates, indicating that incorporation of the dye had no apparent or only negligible effects on plaque morphology. When these images were analyzed in Fiji/ImageJ using Find Maxima (Schindelin, Arganda-Carreras et al. 2012), fluorescent images produced readily detectable, plaque-localized maxima, whereas conventional nonfluorescent images lacked sufficient contrast for similarly reliable automated detection (Figure 1B). Together, these results demonstrate that fluorescent labeling increases plaque contrast sufficiently to render standard plaque assays machine-readable using a simple open-source image-analysis workflow.

**Figure 1.**
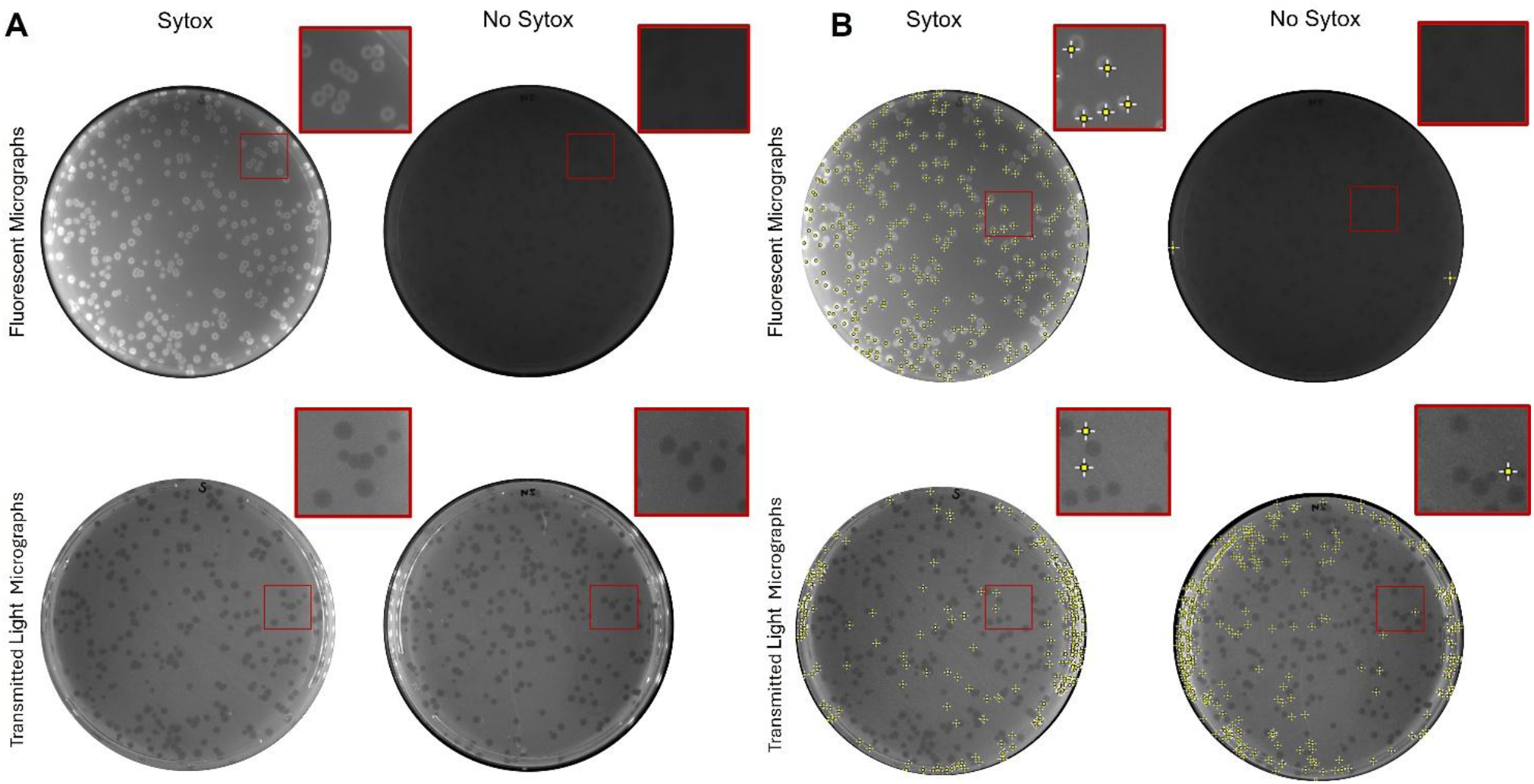
Membrane-impermeant SYTOX dye facilitates software-based plaque identification. (**A**) Fluorescent and transmitted-light images demonstrate that incorporation of SYTOX into the bacterial lawn does not alter T7 plaque formation relative to lawns without SYTOX. (**B)** SYTOX incorporation produces fluorescent plaques for automated plaque counting with the Fiji/ImageJ Find Maxima feature (Schindelin, Arganda-Carreras et al. 2012). Images processed with Fiji/ImageJ: Prominence > 18.00, Strict, Exclude edge maxima, and Output Type: Point Selection.

### 3.2 Fluorescent automated counting works across diverse phage-host systems

We next tested whether this approach extended beyond a single coliphage. Across the systems evaluated, fluorescent plaque labeling supported machine counting for phages with distinct genome types, lifestyles, hosts, and plaque phenotypes (Figure 2A). In addition to T7, the assay enabled accurate machine counting of three additional phages of Gram-negative hosts, while a Gram-positive system is discussed separately below. Key biological features of the phage-host systems examined here are summarized in Table 1.

**Table 1:**
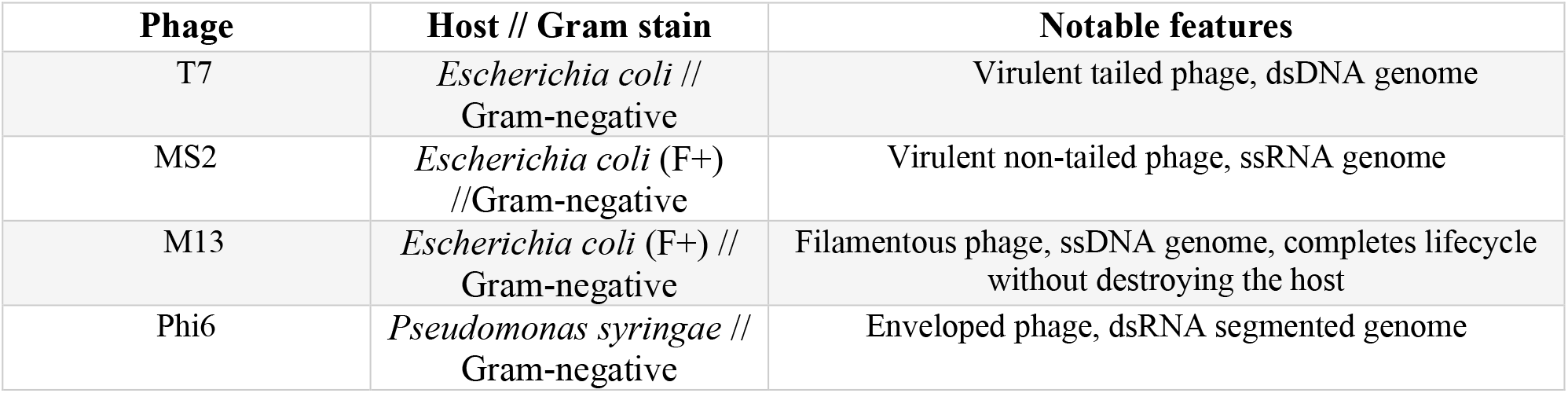
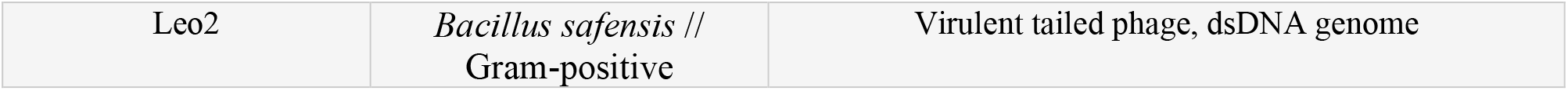
Phage-host systems used to evaluate fluorescent plaque labeling. The panel of phage-host systems was selected to span diverse phage genome types, virion architectures, infection strategies, and host backgrounds, including Gram-negative and Gram-positive bacteria. Features listed here summarize the biological diversity represented in the validation set.

| Phage | Host // Gram stain | Notable features |
| --- | --- | --- |
| T7 | *Escherichia coli* // Gram-negative | Virulent tailed phage, dsDNA genome |
| MS2 | *Escherichia coli* (F+) //Gram-negative | Virulent non-tailed phage, ssRNA genome |
| M13 | *Escherichia coli* (F+) // Gram-negative | Filamentous phage, ssDNA genome, completes lifecycle without destroying the host |
| Phi6 | *Pseudomonas syringae* // Gram-negative | Enveloped phage, dsRNA segmented genome |
| Leo2 | *Bacillus safensis* // Gram-positive | Virulent tailed phage, dsDNA genome |

**Figure 2.**
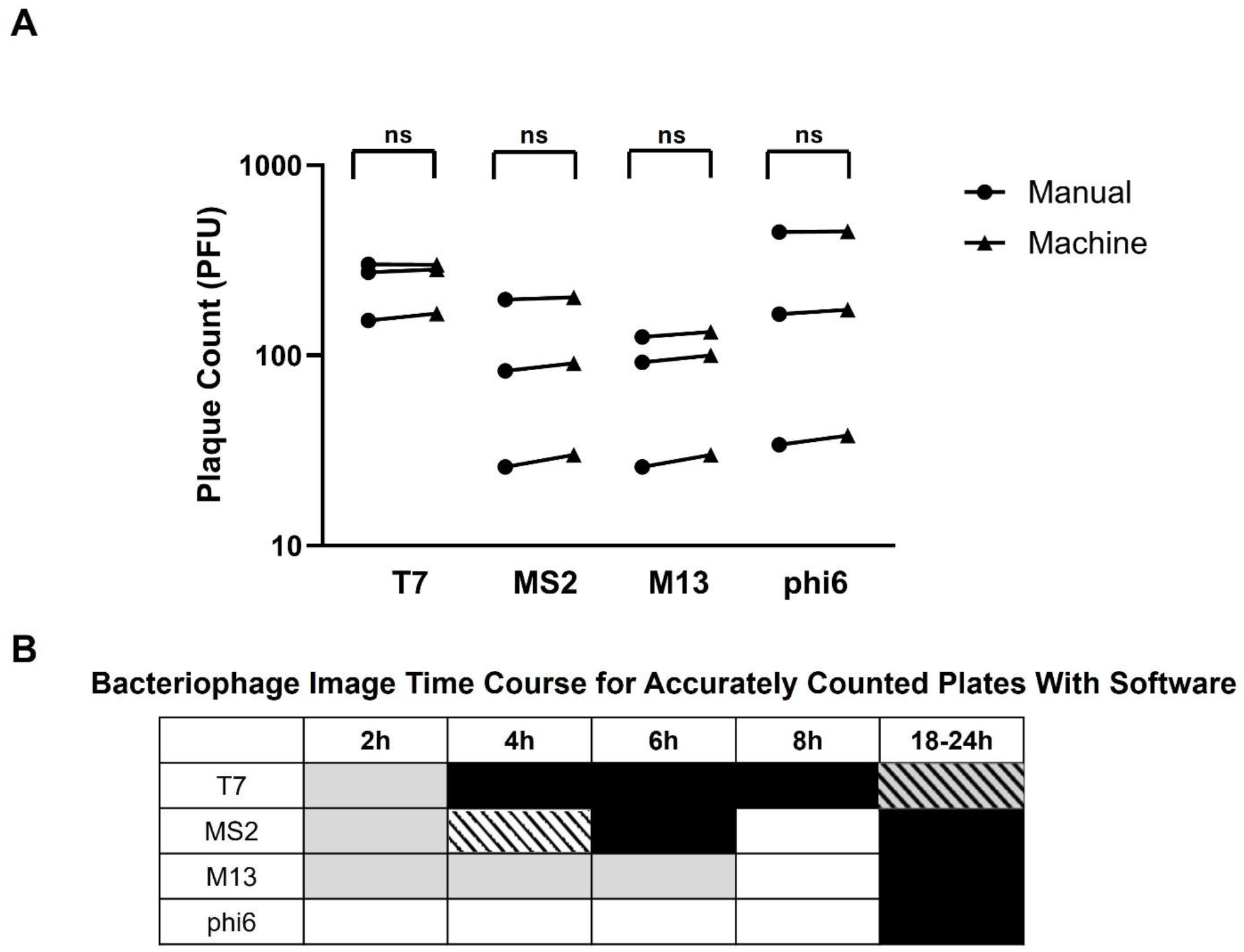
Machine-based counts accurately reflect manual counting in SYTOX-incorporated soft-agar overlays. **(A)** Manual and machine-based (Fiji/ImageJ) plaque counts for T7, MS2, M13, and phi6 from SYTOX-labeled plates. Manual counts were performed at all time points where plaques were visible and confirmed at 24 h for all phages. Automated counts were obtained from fluorescent images acquired at the following timepoints: 6 h for T7 and 24 h for MS2, M13, and phi6. Automated counts were considered equivalent to manual counts when they fell within the expected Poisson counting uncertainty of the manual count, 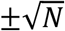. **(B)** Time windows in which fluorescent plaque signal supported machine-based counting within Poisson uncertainty as described above. Black shading indicates emperically determined useful machine-based counting windows, whereas gray shading indicates imaged time windows in which fluorescent plaque signal was insufficient for reliable automated counting under the initial workflow. Gray shading with diagonal black lines denotes conditions that were not initially machine-countable but were later recovered through workflow modification. White shading with diagonal black lines denotes conditions in which machine coutning does not consistently achieve accurate counting within Poisson uncertainty across replicates. Each time window classification was based on two or more independent replicate plates.

M13 provided an informative boundary case. Unlike the other phages examined here, M13 does not lyse its host to release progeny(Marvin 1998, Cahill and Young 2021), and we therefore expected it to be poorly compatible with a dye-based assay that reports nucleic-acid accessibility associated with membrane compromise or cell death. Consistent with this expectation, no useful fluorescent plaque signal was observed in same-day measurements. However, M13 plaques became fluorescent by 24 h and were readily machine-countable at the conventional next-day endpoint. Across all phage systems examined, automated fluorescent counts agreed with manual plaque counts within the expected Poisson counting uncertainty, 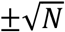, indicating that automated enumeration produced equivalent estimates of plaque abundance (Figure 2). Replicate-level counts across all phages and timepoints are provided in Supplementary Table S1. Together, these systems show that the method is applicable across diverse phage types and its successful use in *E. coli* and *P. syringae* host systems suggest broad portability to Gram-negative plaque assay backgrounds.

### 3.3 Fluorescent labeling can support machine counting during early plaque development

We next explored whether fluorescent labeling could support machine counting as plaques began to develop. Across the systems tested, fluorescent plaques became machine-countable at approximately the same stage that plaques first became visible by eye, with substantially improved contrast for software detection (**Figure 2B)**. Early plaques were often difficult to segment from transmitted-light images of bacterial lawns, whereas corresponding fluorescent images showed discrete puncta or micro-plaques that could be machine-counted and qualitatively linked to countable plaques at later timepoints. These observations indicate that fluorescent labeling can render developing plaques machine-readable during early assay progression. **Figure 2B** summarizes the timepoints and conditions that were most useful for exploratory machine counting across the systems tested. Representative automated counting outputs for these conditions are shown in Supplementary Figures.

T7, which is known to form large plaques, also highlighted that this useful window can close under the initial workflow at later timepoints: by 24 h, plaques had expanded sufficiently that dye signal accumulated at the plaque periphery, producing ring-like labeling that was not well suited to the original counting parameters. However, next-day T7 plaques remained machine-countable after modified handling and adjusted analysis parameters.

Thus, for rapidly expanding plaques such as T7, the method is best paired with same-day imaging or alternative labeling strategies discussed below.

### 3.4 A Gram-positive Leo2 system defines an important compatibility limit and motivates a fallback workflow

We next tested whether the assay was compatible with a Gram-positive phage-host system using Leo2 and *Bacillus safensis* (Badran, Morales et al. 2018, Humphrey, Mackenzie et al. 2023). In contrast to the Gram-negative systems described above, incorporation of dye at the time of plating blocked plaque formation even though the bacterial lawn formed normally (**Figure 3A**). This result identified an important compatibility limit of the pre-incorporation workflow and prompted us to test whether labeling could instead be performed after plaques had formed under standard no-dye conditions. In this post-labeling workflow, SYTOX was applied in buffer to the agar surface after incubation, allowed to absorb and dry, and then imaged. This approach restored fluorescent plaque labeling and enabled accurate machine counting consistent with the performance observed in the compatible pre-incorporation systems.

**Figure 3.**
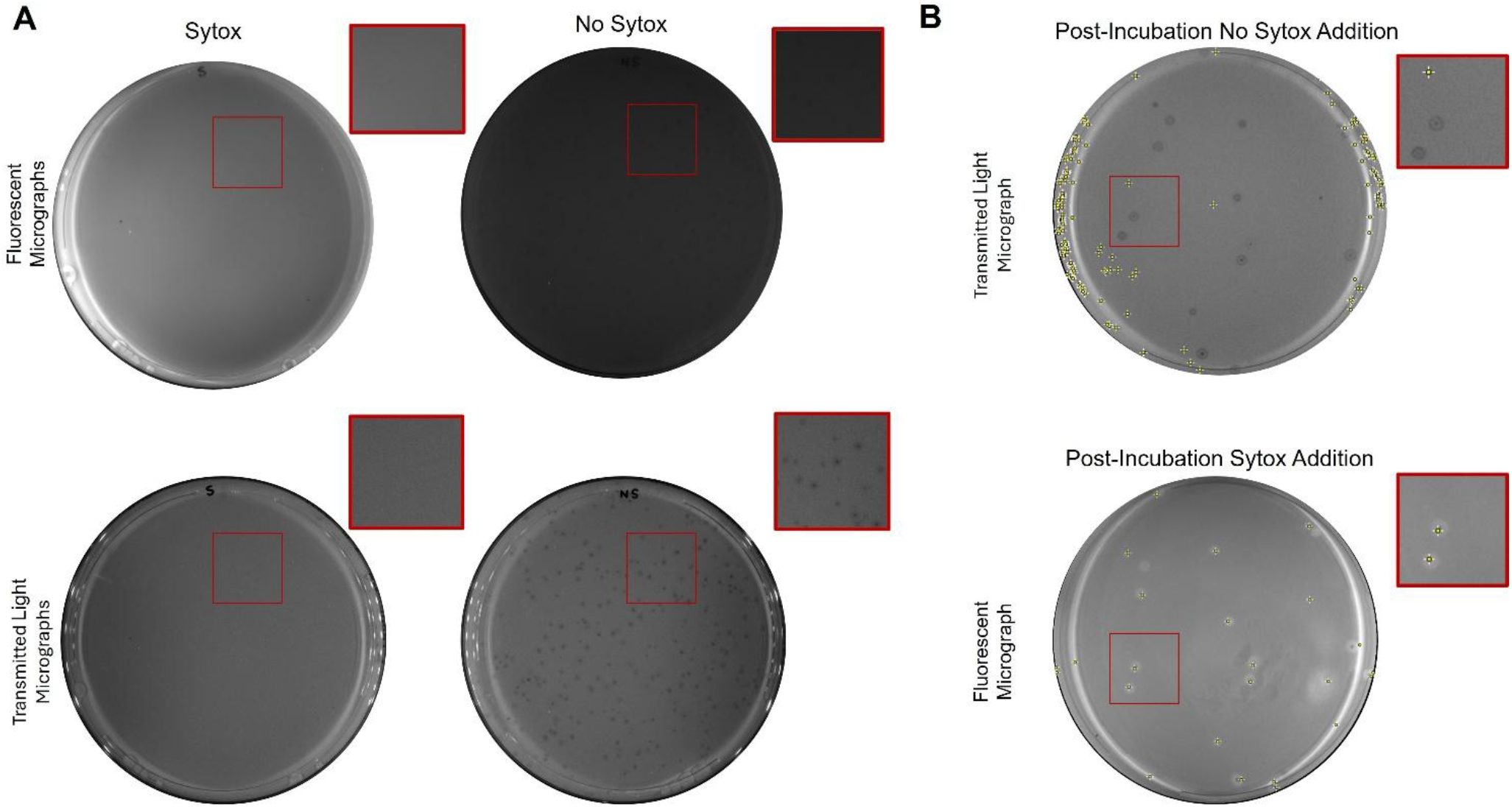
SYTOX dye inhibits Leo2 plaque formation in soft-agar overlay assays. (**A)**. Fluorescent and transmitted-light pictures demonstrate that incorporation of SYTOX into the bacterial lawn inhibits Leo2 plaque formation. (**B)** Transmitted light image demonstrates normal Leo2 plaque formation. Post-incubation addition of SYTOX produces fluorescently labeled plaques that enable machine counting. Images processed with Software: Prominence > 20.00, Strict, Exclude edge maxima, and Output Type: Point Selection.

T7 next-day plaques, which were difficult to count under the initial workflow because of their large size and peripheral dye accumulation, could also be handled using the same general workaround applied to Leo2. When labeled after plaque development and analyzed with adjusted prominence values, next-day T7 plaques remained machine-countable (**Figure 4**).

**Figure 4.**
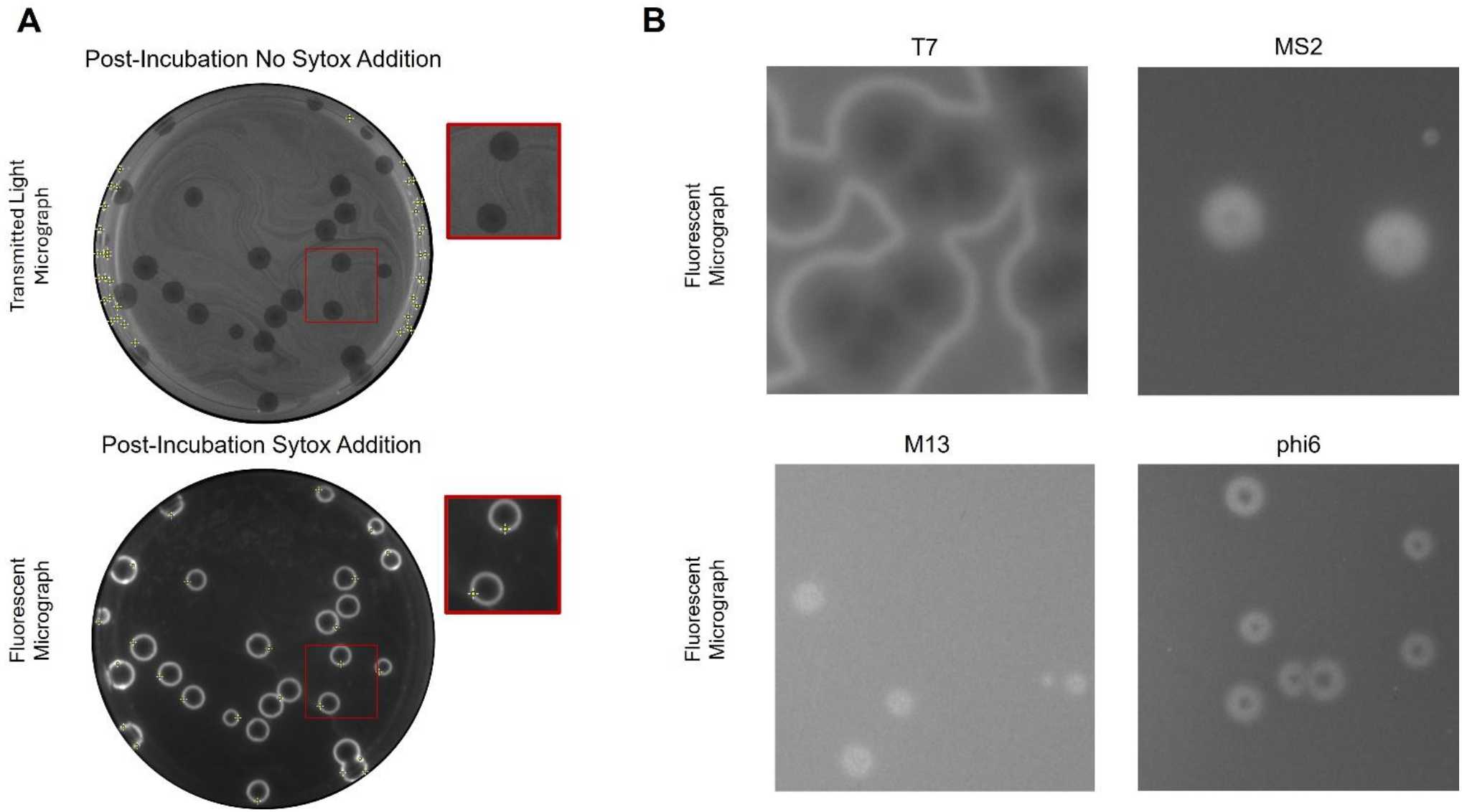
Post-labeling restores machine-countable fluorescence to next-day T7 plaques and highlights phage-dependent plaque morphologies at 24 h. **(A)** Addition of SYTOX after plaque formation produces fluorescently labeled T7 plaques that enable accurate machine counting. Images processed with Software: Prominence > 90.00, Strict, Exclude edge maxima, and Output Type: Point Selection. **(B)** Fluorescent micrographs demonstrate plaque formation and morphological plaque characteristics for T7, MS2, M13, and phi6 at 24 h when SYTOX was incorporated during plating. Each bacteriophage produced distinct fluorescent plaque morphologies.

We next sought to test whether the Leo2 plaque-blocking phenotype observed in agar could be explained by SYTOX interfering with phage interaction with *B. safensis*. We measured supernatant-recoverable Leo2 in host-containing and host-free liquid reactions with and without dye. In no-cell broth controls, Leo2 titers recovered from clarified supernatants remained at ∼1x10^9^ PFU/mL at both 10 and 30 min regardless of whether SYTOX was present (**Figure 5 compare red and black traces**). This indicates that the dye did not measurably reduce free-phage recovery under these conditions. In contrast, addition of *B. safensis* cells markedly reduced the recovery of Leo2 ∼100-fold relative to no-cell controls by the first sampled timepoint (**Figure 5**). The reduced recovery is consistent with a majority of free phage adsorbing to the host during the infection step. Recovered titers remained at similar levels at 30 min, indicating that most of the host-dependent depletion of recovered phage occurred within the first 10 min of the assay. Under this liquid-phase design, any substantial SYTOX-dependent effect on early free-phage interaction would therefore likely have been evident by the sampled timepoints.

**Figure 5.**
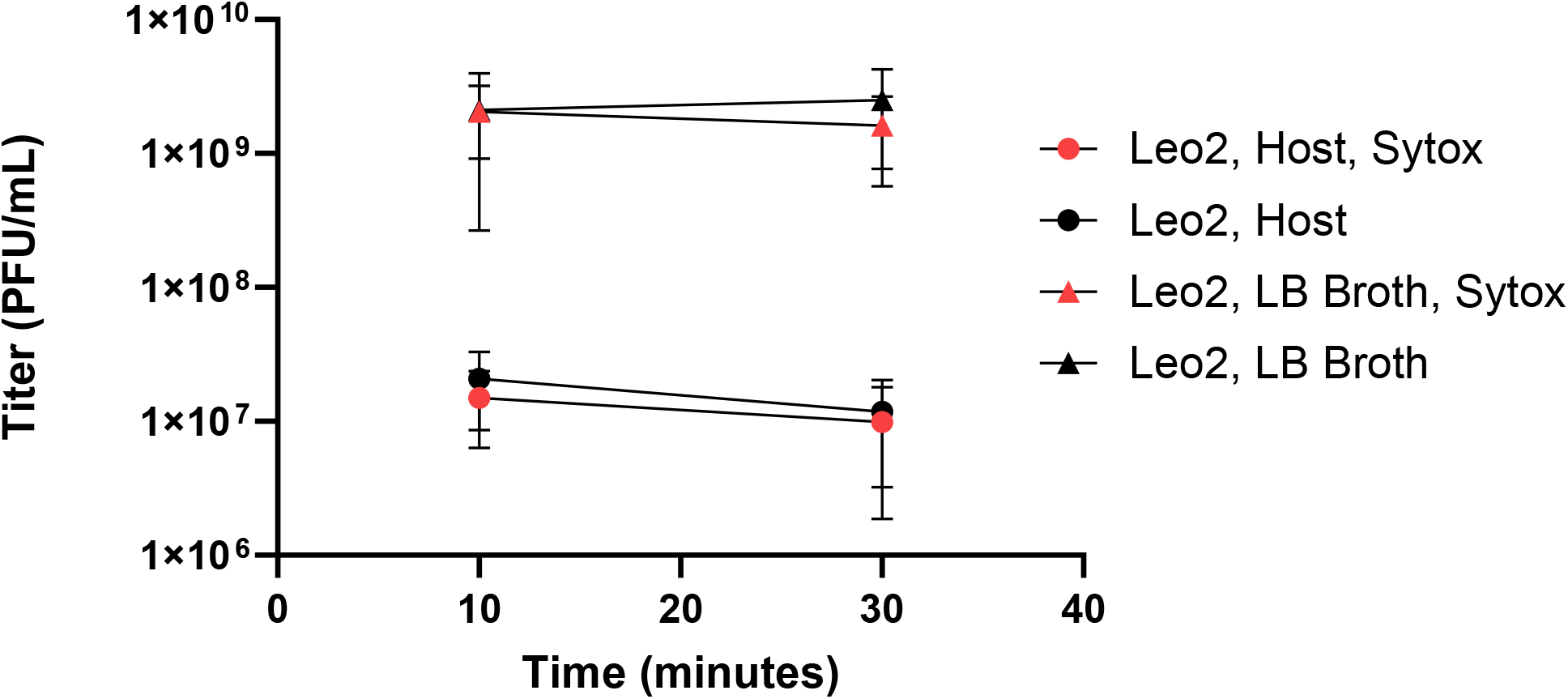
Liquid-phase Leo2 recovery assay in the presence and absence of host cells and SYTOX. Time zero indicates addition of *Bacillus safensis* host cells to mixtures containing Leo2 phage in LB broth with or without SYTOX. Supernatant-recoverable phage was measured from clarified samples collected at 10 and 30 min. At both 10 and 30 min, host-containing reactions had significantly lower recovered Leo2 than the corresponding no-cell broth controls, both in the presence and absence of SYTOX. No significant differences were detected between SYTOX-treated and no-dye conditions within either host-containing or host-free samples, and no significant differences were observed between 10 and 30 min within any condition. Values were analyzed as log_10_-transformed PFU/mL. Markers represent mean, and the error bars represent standard deviation for three independent replicates per condition.

Statistical analysis of log_10_-transformed PFU/mL values supported this interpretation. Three-way ANOVA identified host-cell presence as the only significant factor (p < 0.0001), whereas SYTOX presence, time, and all interaction terms were not significant. Planned Šídák-corrected comparisons showed significant host-dependent depletion of supernatant-recoverable Leo2 at both timepoints but no significant difference between SYTOX-treated and no-dye reactions in either host-containing or host-free conditions. No significant differences were observed between 10 and 30 min within any condition. Together, these results indicate that, under the tested liquid conditions, SYTOX did not measurably alter Leo2 recovery either in broth alone or in the presence of host cells.

### 3.5 Dye labeling generally preserves recovered phage yield for downstream workflow

A final practical question was whether fluorescent plaque labeling remained compatible with downstream phage recovery workflows. In many plaque-based experiments, enumeration is only one step, and plaques are subsequently recovered for propagation, purification, or further characterization. This consideration is especially relevant for future automation-oriented workflows in which plaque detection, quantification, and recovery may be linked within a single pipeline. It was also made more important by the Leo2 result above, where dye present at plating blocked plaque formation and raised the possibility that prolonged dye exposure might compromise recoverable phage infectivity even when plaques were successfully formed.

To test this, plaques were recovered by a standard pickate method in which individual plaques were cored from the agar, resuspended in buffer, and then re-titered on fresh indicator lawns. Under these conditions, MS2 showed a statistically significant reduction in recoverable infectivity after dye exposure, whereas Leo2 did not (**Figure 6**). Notably, the reduction observed for MS2 did not measurably impair plaque formation in the corresponding overlay assay. Likewise, post-labeled Leo2 plaques yielded recoverable viable phage from independent pickates with no detectable loss relative to controls.

**Figure 6.**
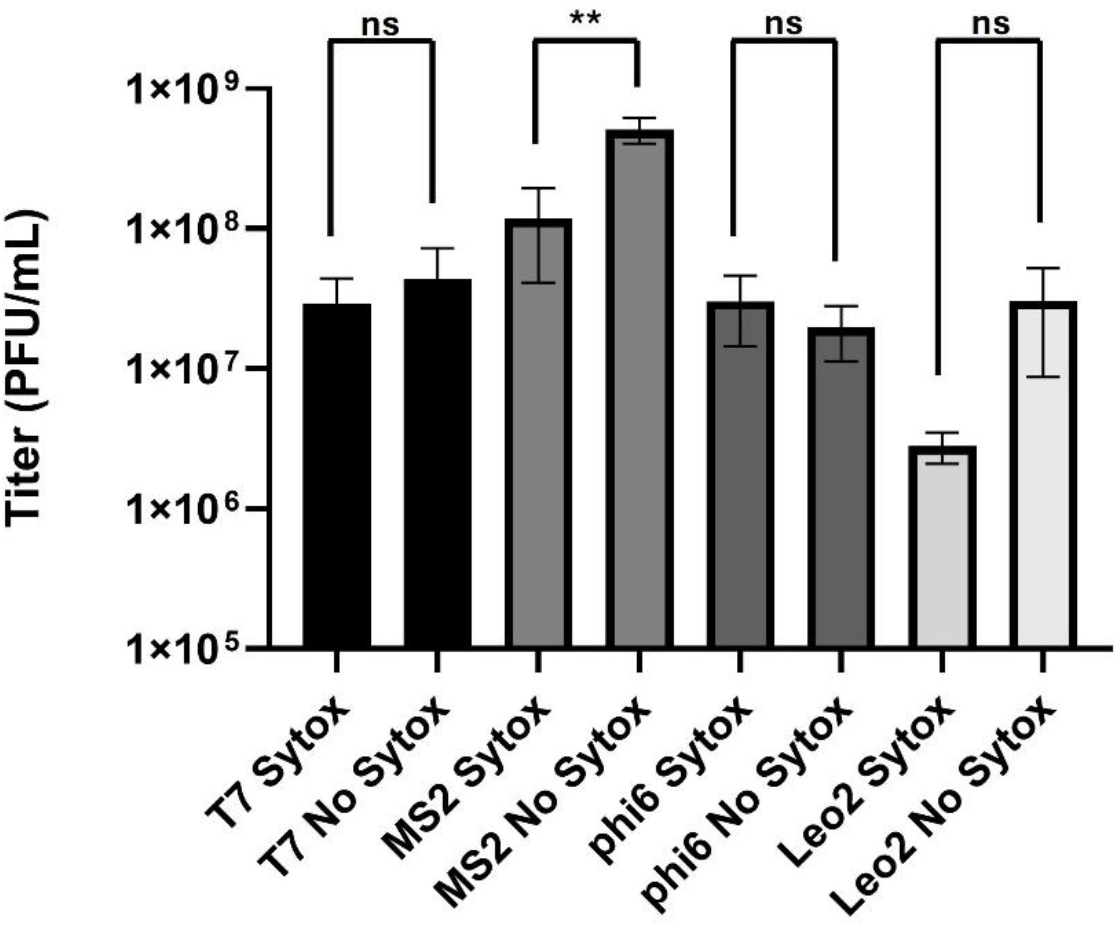
Plaque Titer for T7, MS2, phi6, and Leo2. Individual plaques picked from plates containing SYTOX or plates containing no addition of SYTOX. T7, MS2, and phi6 plaques were picked from plates in which SYTOX was incorporated in the bacterial host lawn. Leo2 plaques were picked from plates prepared with an alternative SYTOX addition method. To test for statistical significance, an unpaired t-test was performed, where **** = P< 0.0005, *** = P< 0.005,** = P < 0.05, * = P< 0.5 in which n = 3 independent replicates.

Taken together, the liquid-phase interaction assay and plaque-recovery experiment do not identify the precise mechanism by which SYTOX blocks Leo2 plaque formation in agar, but they argue against the simplest explanations. SYTOX did not measurably reduce free Leo2 recovery in host-free broth controls, and post-labeled Leo2 plaques yielded recoverable viable phage from agar cores at levels not significantly different from no-dye controls, arguing against direct dye-mediated phage damage as a sufficient explanation for the Leo2 plaque-blocking phenotype. In addition, SYTOX did not measurably alter host-dependent depletion of free phage from clarified supernatants, arguing against a large effect on early liquid-phase phage-host association under the tested conditions. The remaining incompatibility therefore appears specific to plaque establishment in the agar-overlay context. These findings further indicate that, in compatible workflows, fluorescent labeling can be incorporated into plaque assays without generally preventing downstream plaque recovery and propagation.

### 3.6 Fluorescent labeling improves plaque detection capabilities of large language models

Given the growing use of multimodal large language models (LLMs) in image interpretation tasks, we asked whether SYTOX also improves plaque counting in an LLM-based workflow. To investigate LLM-based plaque counting, we directly compared counts of captured images with and without SYTOX using a multimodal LLM (Claude AI, Anthropic) to the methodology we described above. As a representative plate, we used the SYTOX-containing T7 plate at 6 h (**Figure 1A**). A manual count was used as the ground truth for the number of plaques within a bounded region of the plate, with the same region subsequently used for all counting methods (**Figure 7A**). Transmitted-light and fluorescence images of this SYTOX-containing plate were then analyzed by the LLM (see Methods) or by Fiji/ImageJ using Find Maxima (Schindelin, Arganda-Carreras et al. 2012)(**Figure 7B and C**). Both the LLM and Find Maxima undercounted plaques in transmitted-light images, in which SYTOX was present but fluorescence was not collected. In contrast, fluorescence imaging provided a detectable signal that improved plaque identification. The deviations of the transmitted-light counts from the manual ground truth exceeded the expected √N variation associated with a Poisson counting process, indicating that the differences were unlikely to be explained by counting noise alone.

**Figure 7.**
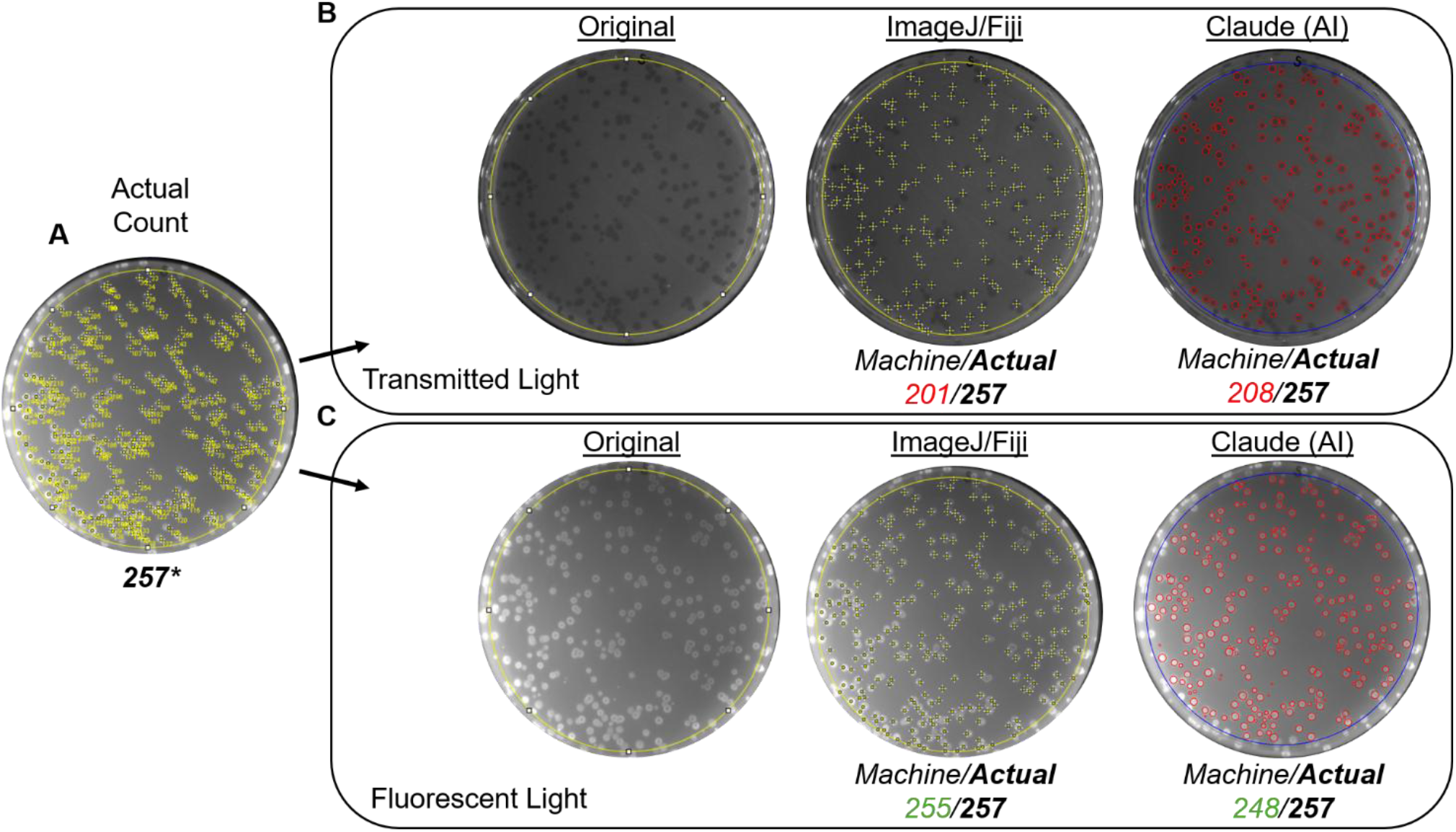
Comparison of plaque counting methodology. (**A)** A fluorescent light micrograph following SYTOX dye incorporation into the bacterial imaged for T7 plaque formation at 6 h was manually counted using ImageJ/Fiji multi-selection tool, which applies numbers to hand-counted zones. This is used to generate a “ground truth” count of 257 plaques. The yellow circle represents a boundary zone applied by the LLM (blue in images to the right) so that comparisons remain consistent across groups. **(B)** Transmitted light micrographs (not exciting SYTOX fluorophore) were counted using both the ImageJ/Fiji “Find maxima” feature, with image inversion, or a large language model (LLM; Claude AI, Anthropic) to assess plaque counts. **(C)** Fluorescent light micrographs (which excite the SYTOX fluorophore) were similarly counted to compare the performance of machine-based counting tools under different light conditions for plates containing SYTOX dye. All counts are shown in relation to actual plaque counts (black, bolded), where red coloed font indicates counts that deviate by more than the Poisson uncertainty (square root of the number of plaques manually counted). Green color font indicates machine-based counts that fall within the square root of the number of plaques manually counted.

However, when fluorescent light images were used to excite and detect the SYTOX fluorophore, both the LLM and Find Maxima methodologies performed similarly, with plaque counts falling within Poisson counting uncertainty of the manual count. Together, these data indicate that a frontline multimodal LLM does not provide facile plaque counting from standard transmitted-light plate images. Rather, both LLM-based and conventional image-analysis approaches benefit substantially from the use of SYTOX as a plaque contrast agent, as introduced here for phage plaque visualization.

## 4 Discussion

### 4.1 A practical route to automated plaque counting

The primary contribution of this study is a simple route for converting standard bacteriophage plaque assays into a machine-readable format for automated counting. Rather than relying on reporter phages, custom machine-learning workflows, or specialized imaging systems, the method uses a membrane-impermeant nucleic acid dye, standard soft-agar plaque workflows, and an open-source Fiji/ImageJ analysis pipeline based on Find Maxima. The key result here is that contrast enhancement by SYTOX labeling improves the image input itself, making automated counting feasible with simple, widely available software while also benefiting AI-based approaches (discussed below). Across the compatible systems tested, automated fluorescent counts agreed with manual counts within the inherent Poisson uncertainty of plaque counting, indicating that the automated workflow produced equivalent estimates of plaque abundance. In this sense, the method does not aim to outperform human counters in absolute terms, but to render plaque assays sufficiently high-contrast that simple software can perform the counting task reliably. This practical accessibility may also extend across imaging platforms. In addition to the Syngene G:BOX system used for the main experiments, we observed comparable machine-countable SYTOX plaque contrast using a standard blue-light transilluminator (Supplementary Figure S7). Although we did not systematically benchmark multiple instruments, this result suggests that the method is not tightly coupled to a single imaging system.

We also tested whether a representative state-of-the-art large language model (Claude)(Anthropic 2026) could count plaques from the same images used in the non-AI workflow. This comparison is informative because the T7 system represents a relatively undemanding test case for automated plaque counting: plaques are large, clear, and morphologically regular, and should therefore be among the easier conventional plaque images for automated interpretation. Even in this favorable setting, both Claude and Fiji/Find Maxima (Schindelin, Arganda-Carreras et al. 2012)undercounted conventionally imaged transmitted-light plates (**Figure 7B**), whereas SYTOX fluorescence brought both workflows within Poisson counting uncertainty of the manual count (**Figure 7C**). This indicates that the principal advantage of the dye is not tied to one software ecosystem, but to improved contrast at the level of the assay image itself. In that sense, SYTOX labeling appears complementary to both conventional image-analysis workflows and future AI-assisted approaches.

We chose a web-based multimodal LLM tool that incorporates VLM capabilities, as they are widely distributed and require no special system to run. Future efforts in benchmarking to a dedicated VLM trained on bacteriophage plaques such as OnePetri (Shamash and Maurice 2021) are expected to show similar performance enhancements as the underlying methodologies are the same.

At the same time, the present workflow should be viewed as a calibrated semi-automated approach rather than a universal out-of-box replacement for all plaque-counting applications. As with other plate-image analysis methods, performance can be influenced by factors such as plaque morphology, plaque density, illumination uniformity, edge glare, cropping, and the choice of detection parameters such as prominence. The present study was designed to establish the value of SYTOX as a contrast reagent and to demonstrate practical machine readability across representative systems, rather than to validate a fully blinded, operator-independent workflow across all plaque types and imaging conditions. In practice, we expect the method to be most useful in high-volume experimental settings where plaque biology, imaging geometry, and analysis settings can be calibrated once and then applied repeatedly across many plates. Under those conditions, the up-front tuning effort is likely to pay off substantially by reducing manual counting burden. Where such fine-tuning is impractical, AI-assisted workflows may ultimately provide a more general route to automation, but the results here show that SYTOX improves the image input for both AI-based and LLM-free analysis strategies.

### 4.2 Fluorescent labeling reveals biologically distinct patterns of plaque development

The timing and morphology of fluorescent plaque labeling depended strongly on phage biology. In lytic systems, fluorescence emerged during plaque development and frequently rendered plaques machine-countable as soon as they became meaningfully visible by eye. M13, by contrast, did not yield useful same-day fluorescence and became detectably fluorescent only at the conventional next-day endpoint. This difference is biologically informative. The canonical description of M13 infection is based largely on liquid broth culture, where infected cells remain viable while continuously extruding progeny virions. Plaque formation in agar presents a markedly different physical environment. Within a spatially constrained gel matrix, chronically infected cells remain locally confined while continuously producing phage under conditions of prolonged infection and progressive nutrient limitation. Under these conditions, the physiological fate of infected cells may differ substantially from that observed in well-mixed liquid culture, ultimately leading enough cells within mature plaques to lose membrane integrity and permit SYTOX labeling. If this timing difference proves generalizable, delayed fluorescence may provide a simple phenotypic clue for distinguishing filamentous chronic phages from virulent lytic phages during phage discovery or characterization.

Differences were also evident in the spatial distribution of fluorescence during plaque maturation. Developing plaques of all lytic phages (i.e., excluding M13) examined initially appeared as uniformly fluorescent puncta; however, as plaques matured, fluorescence progressively redistributed toward the plaque perimeter (**Figure 4B**). This effect was subtle in the relatively small plaques produced by MS2 and M13, more apparent in phi6, and most pronounced in the large plaques formed by T7, which frequently developed distinct fluorescent rings. These observations suggest that peripheral redistribution of fluorescence is a general consequence of plaque maturation rather than a phenomenon unique to T7. A simple explanation is that newly lysed cells at the advancing plaque margin continuously release fresh nucleic acids available for SYTOX labeling, whereas nucleic acids within older plaque regions are progressively lost through degradation and diffusion over time. The magnitude of this redistribution likely depends on plaque architecture, plaque size, and the duration over which nucleic acids remain within the plaque environment. T7 may further accentuate this process because it encodes nucleases involved in extensive host chromosomal DNA degradation, recycling host nucleotides for phage DNA synthesis (Mitsunobu, Zhu et al. 2014). However, the present study does not distinguish the relative contributions of phage-encoded nuclease activity, host nucleases released during lysis, diffusion, or other aspects of plaque development to the observed ring morphology.

Together, these observations indicate that fluorescent plaque labeling is more than a contrast-enhancement technique. The timing and morphology of fluorescence reflects the underlying biology and spatial progression of infection within individual plaque systems. Future studies could determine whether differences in fluorescent signal can be exploited as simple phenotypic indicators of phage types or infection strategies.

### 4.3 Compatibility limits and fallback workflows

The Leo2–*Bacillus safensis* system showed that the method is not universal in the simple agar-seeded workflow used at plating. Dye present at the time of plating blocked Leo2 plaque formation even though the host lawn developed normally. This result defined an important compatibility limit and motivated a useful fallback strategy. When plaques were first formed under standard no-dye conditions and labeled afterward, fluorescent plaque detection and machine counting were restored (**Figure 3**). Follow-up experiments narrowed but did not fully explain the basis of this incompatibility. Post-labeled Leo2 plaques yielded recoverable viable phage from agar cores at levels not significantly different from no-dye controls, arguing against direct dye-mediated phage damage as a sufficient explanation for the plaque-blocking phenotype (**Figure 6**). In addition, the liquid-phase interaction assay showed that SYTOX did not measurably reduce free Leo2 recovery in host-free broth controls and did not measurably alter host-dependent depletion of free phage from clarified supernatants (**Figure 5**). Thus, the incompatibility does not appear to arise from simple direct phage inactivation or from a large defect in liquid-phase host association under the tested conditions.

Instead, it appears to depend on features of plaque establishment in semi-solid media that were not recapitulated in the liquid assay. Possible explanations include interference with genome injection, or disruption of lysis-related functions. This is practically important because future researchers may encounter similar system-specific limitations, and post-labeling provides a path forward for machine counting when agar-seeded dye incorporation fails.

From a practical perspective, these observations also define the optimal operating window for fluorescence-assisted plaque enumeration. Large-plaque phages such as T7 were most readily quantified during early plaque development before extensive peripheral redistribution occurred, whereas smaller-plaque systems remained accurately countable even at conventional overnight endpoints. Thus, the assay is optimized not for imaging large plaques formed a day after plating, but for accelerating plaque visualization and improving automated enumeration during the earliest stages of plaque development. Where next-day sampling is unavoidable, future users of the method may be able to improve handling of large-plaque formers by tuning plaque-size determinants such as incubation temperature, inoculum density, or agar concentration to limit late-stage plaque expansion.

### 4.4 Opportunities for higher-throughput phage biology

The broader significance of this work is that it lowers the practical barrier to larger and more quantitative plaque-based experiments. By reducing the manual burden of plaque counting, the method can support more replicate plates, more plaque counts per plate, and therefore stronger statistical confidence in phage quantification experiments. In the near term, we expect the method to be most useful in repeated high-volume workflows where plaque morphology, imaging conditions, and analysis settings can be standardized well enough that the initial calibration effort is repaid across many plates and experiments. The finding that fluorescent labeling is generally compatible with downstream plaque recovery also increases its usefulness for workflows in which enumeration is followed by propagation, purification, or further characterization. Looking forward, this kind of machine-readable plaque assay could serve as a component of more integrated automated or robotic phage workflows, including pipelines in which plaques are detected, counted, recovered, and advanced with limited human intervention. The observed improvement in plaque-image contrast may also make these assays better inputs for future AI-assisted image analysis, even though the present study emphasizes a practical non-AI route to automation. The framework presented here may also help other investigators evaluate dye compatibility in their own phage-host systems and refine the method further. In particular, future improvements in imaging sensitivity, segmentation, and automation may allow reliable counting of discrete early puncta before conventional visual plaque appearance. If that becomes robust, it could increase usable plaque densities per plate well beyond current manual practice and lower the practical barrier to statistically rigorous phage experiments.

## 5 Permission to reuse and Copyright

This is an open-access article distributed under the terms of the Creative Commons Attribution License (CC BY). The use, distribution or reproduction in other forums is permitted, provided the original author(s) and the copyright owner(s) are credited and that the original publication in this journal is cited, in accordance with accepted academic practice. No use, distribution or reproduction is permitted which does not comply with these terms.

## 7 Disclaimer

This paper describes objective technical results and analysis. Any subjective views or opinions that might be expressed in the paper do not necessarily represent the views of the U.S. Department of Energy or the United States Government.

## 9 Author Contributions

AW: Investigation, Data curation, Visualization, Resources, Writing-review & editing

JA: Investigation, Methodology, Software, Visualization, Writing-original draft, Writing-review & editing

DB: Investigation, Funding acquisition, Writing-original draft, Writing-review & editing

JC: Investigation, Conceptualization, Methodology, Project administration, Resources, Supervision, Validation, Visualization, Writing—original draft, Writing—review and editing

## Acknowledgments

We thank the Cahill Laboratory members and the Sandia National Laboratories’ Environmental Systems Biology and Molecular and Microbiology Department for their valuable input during this study. We thank Eric Small for reviewing a pre-submission manuscript draft, suggesting edits, and providing technical feedback. SandiaAI Chat, a version of OpenAI’s GPT-5.4 architecture, was used to assist with ideation, organization, drafting, and proofreading. All authors reviewed, approved, and take responsibility for the final manuscript.

## Data Availability Statement

All datasets generated for this study will be made available in supplementary material.

