## Supplemental Tables (1-3) and Figures (1-7) for "A membrane-impermeant nucleic acid dye converts bacteriophage plaque assays into a machine-readable format for automated counting"

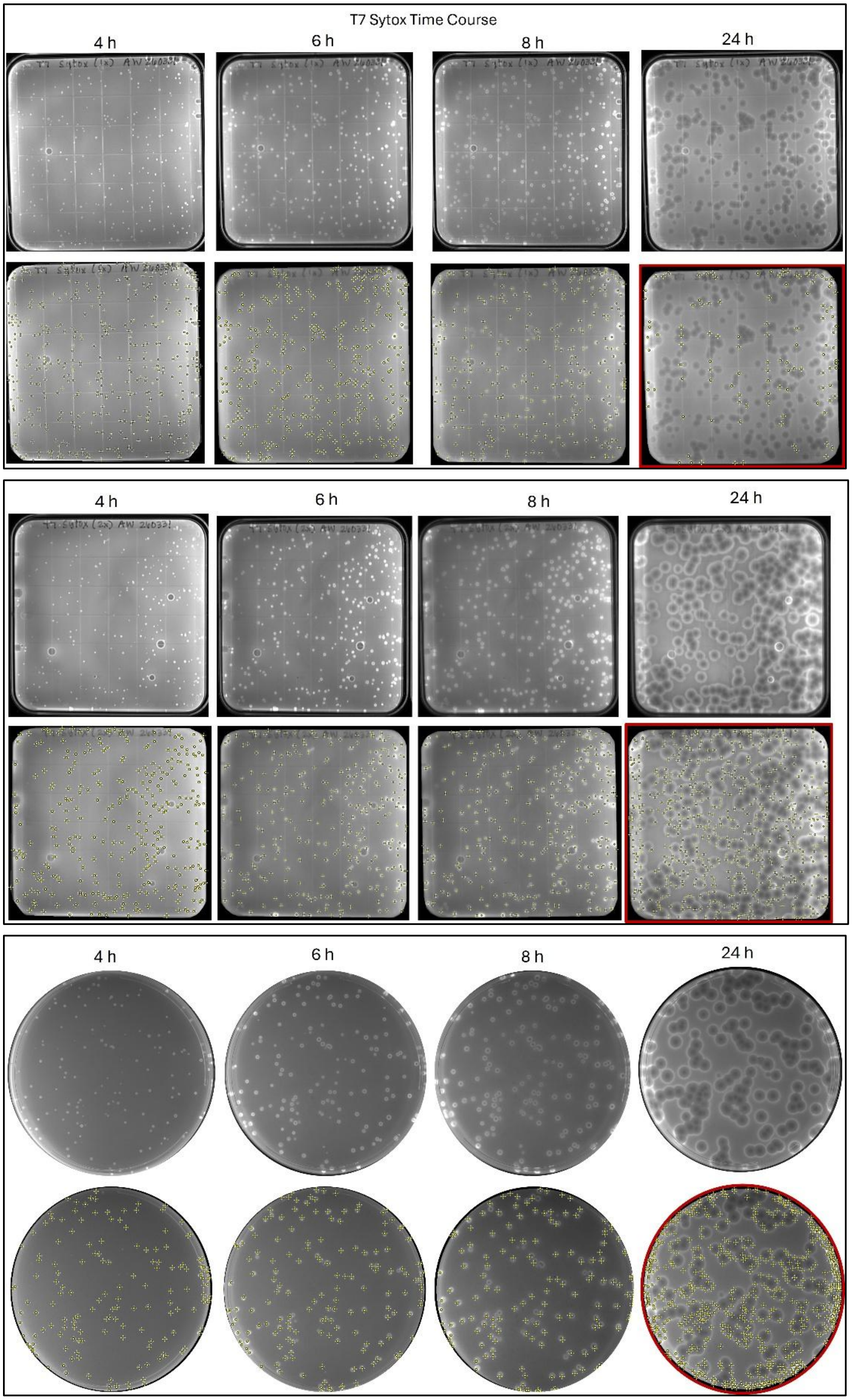

**Figure S1. Machine-based point selections of T7 plaques over time.**

Fluorescent micrographs of T7 plaques demonstrate point selections during machine-based counting at 4, 6, and 8 h, as shown in Figure 2. By 24 h, T7 plaques could not be accurately counted using the same workflow and are outlined in red to indicate inaccurate point selections. As shown in Figure 4, a fallback workflow was implemented at 24 h to enable accurate machine counting of fluorescent T7 plaques.

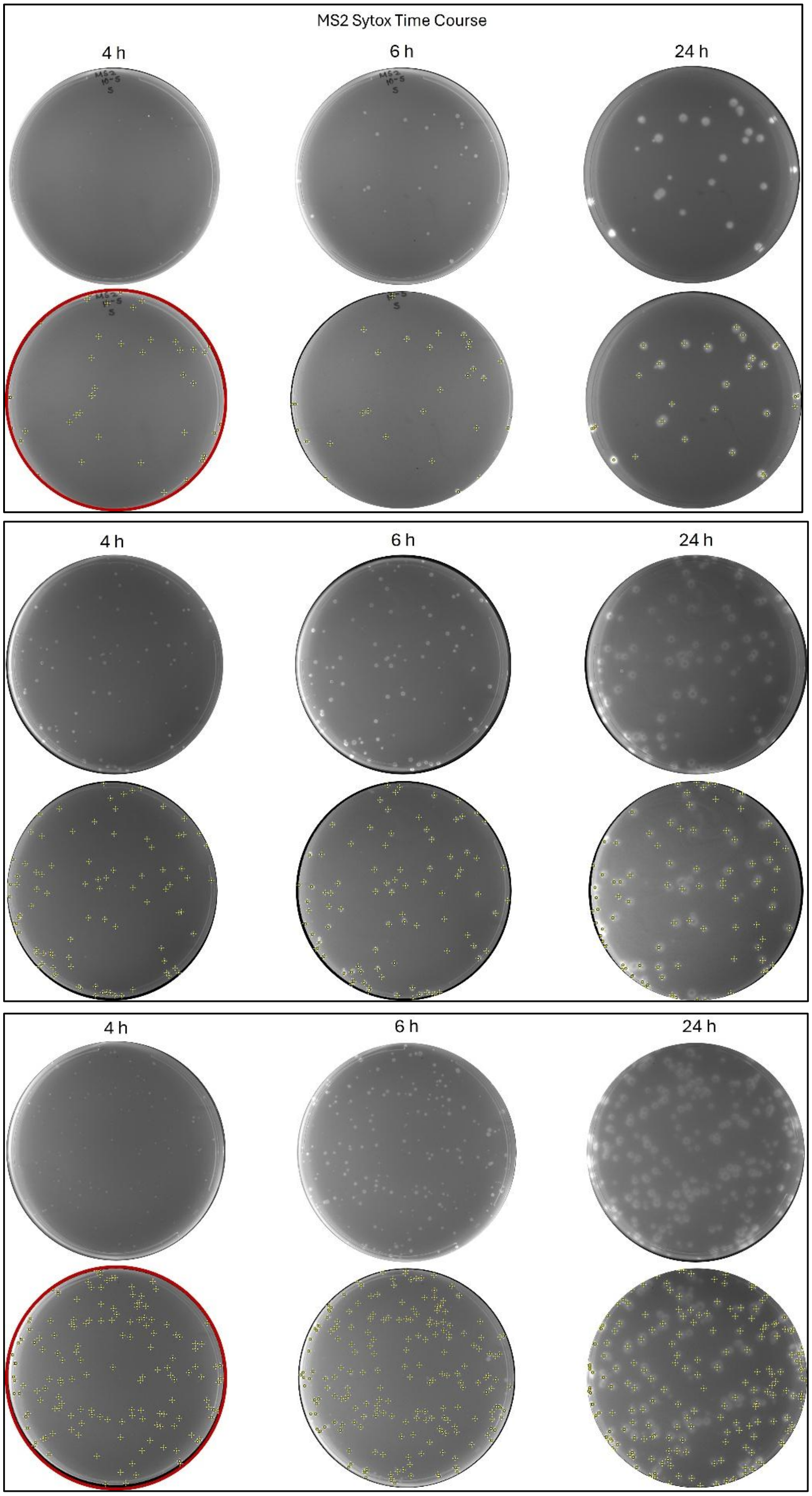

**Figure S2. Machine-based point selections of MS2 plaques over time.**

Fluorescent micrographs of MS2 plaques demonstrate point selections during machine-based counting at 6 and 24 h, as shown in Figure 2. At 4h, conditions do not consistently achieve accurate counting within Poisson uncertainty across replicates and are outlined in red.

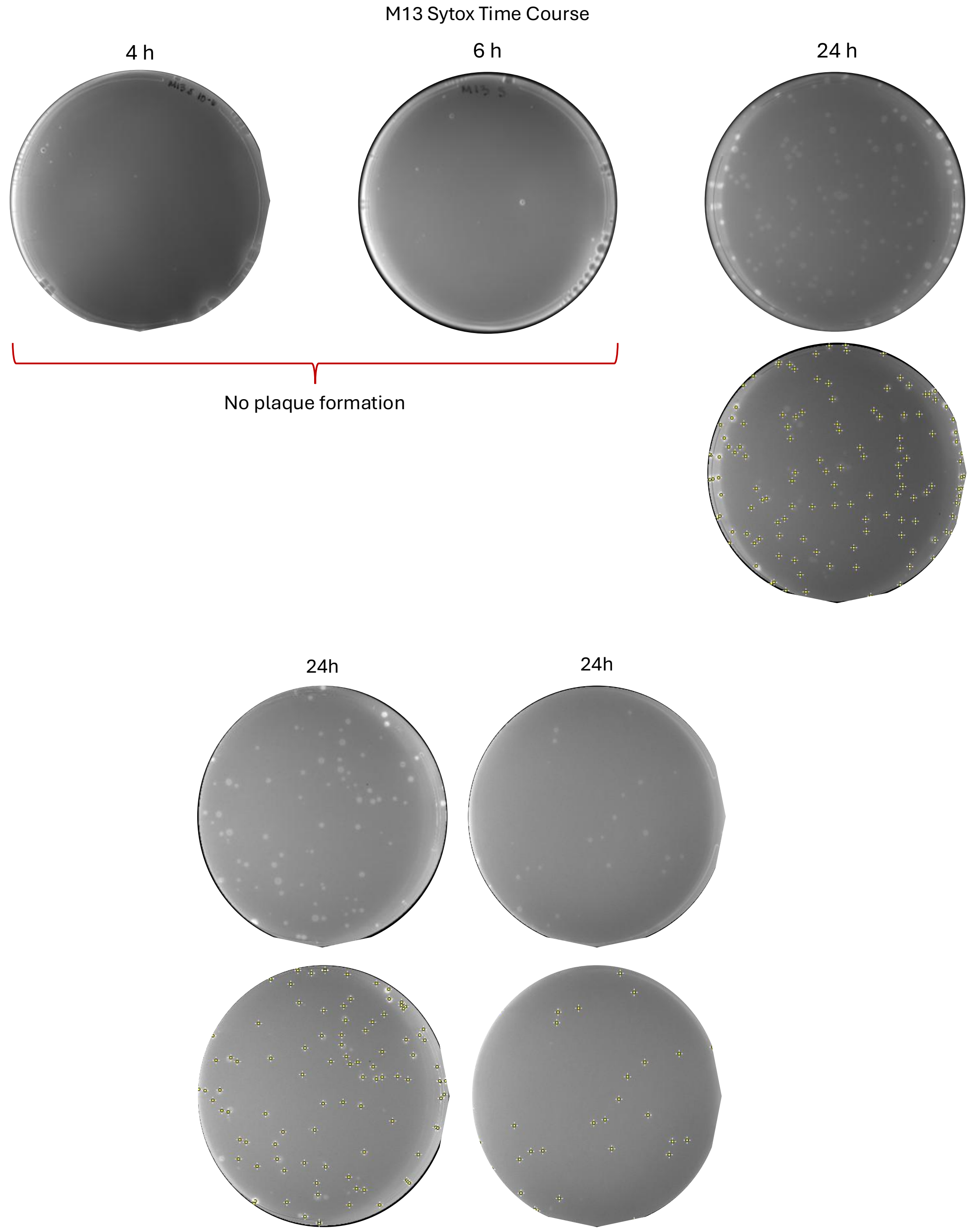

**Figure S3. Machine-based point selections of M13 over time.**

Fluorescent micrographs for M13 display accurate point selections with machine-based counting at 24h for n=3 independent replicates, as shown in Figure 2. As plaques appear outside normal working hours, machine counting with other earlier time points was not investigated.

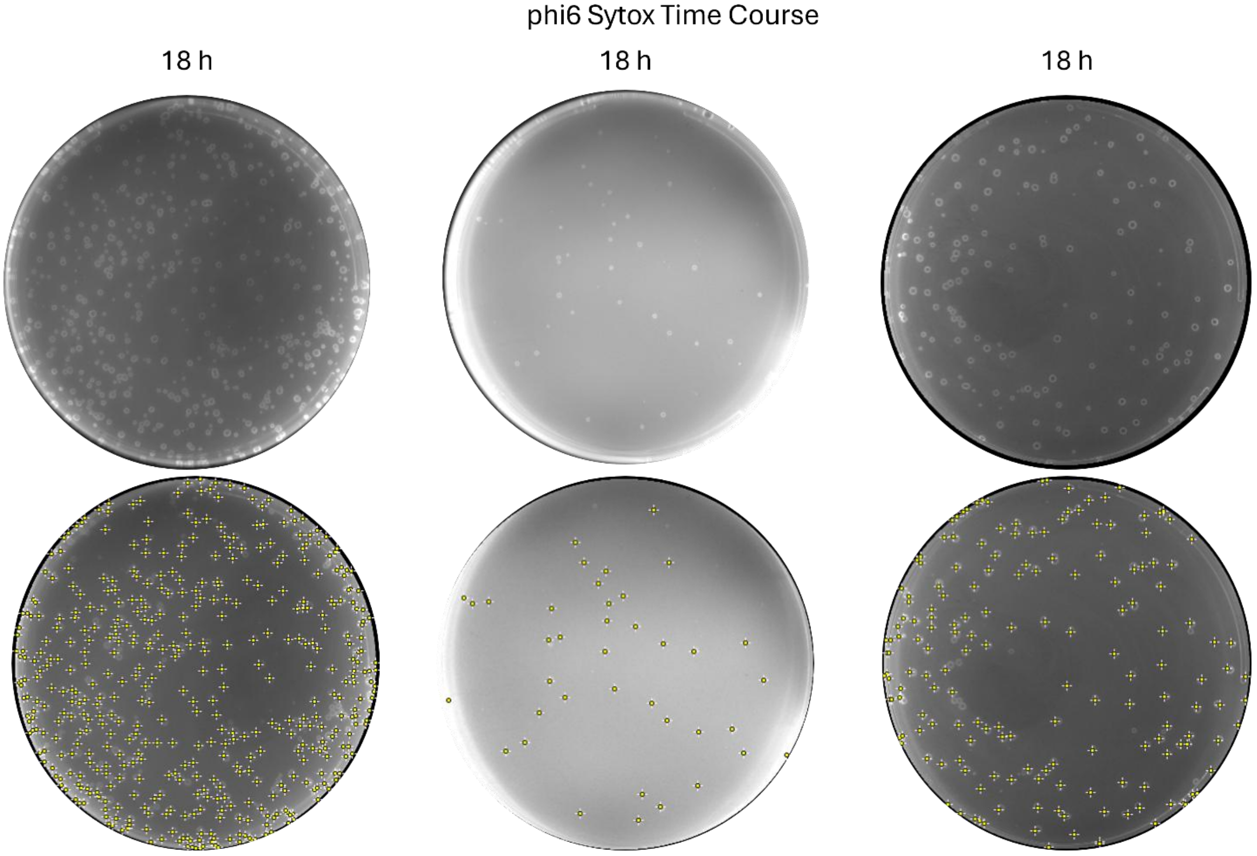

**Figure S4. Machine-based point selections of phi6 plaques over time.**

Fluorescent micrographs for phi6 display accurate point selections with machine-based counting with next day images for n=3 independent replicates, as shown in Figure 2. As plaques appear outside normal working hours, machine counting with earlier time points was not investigated.

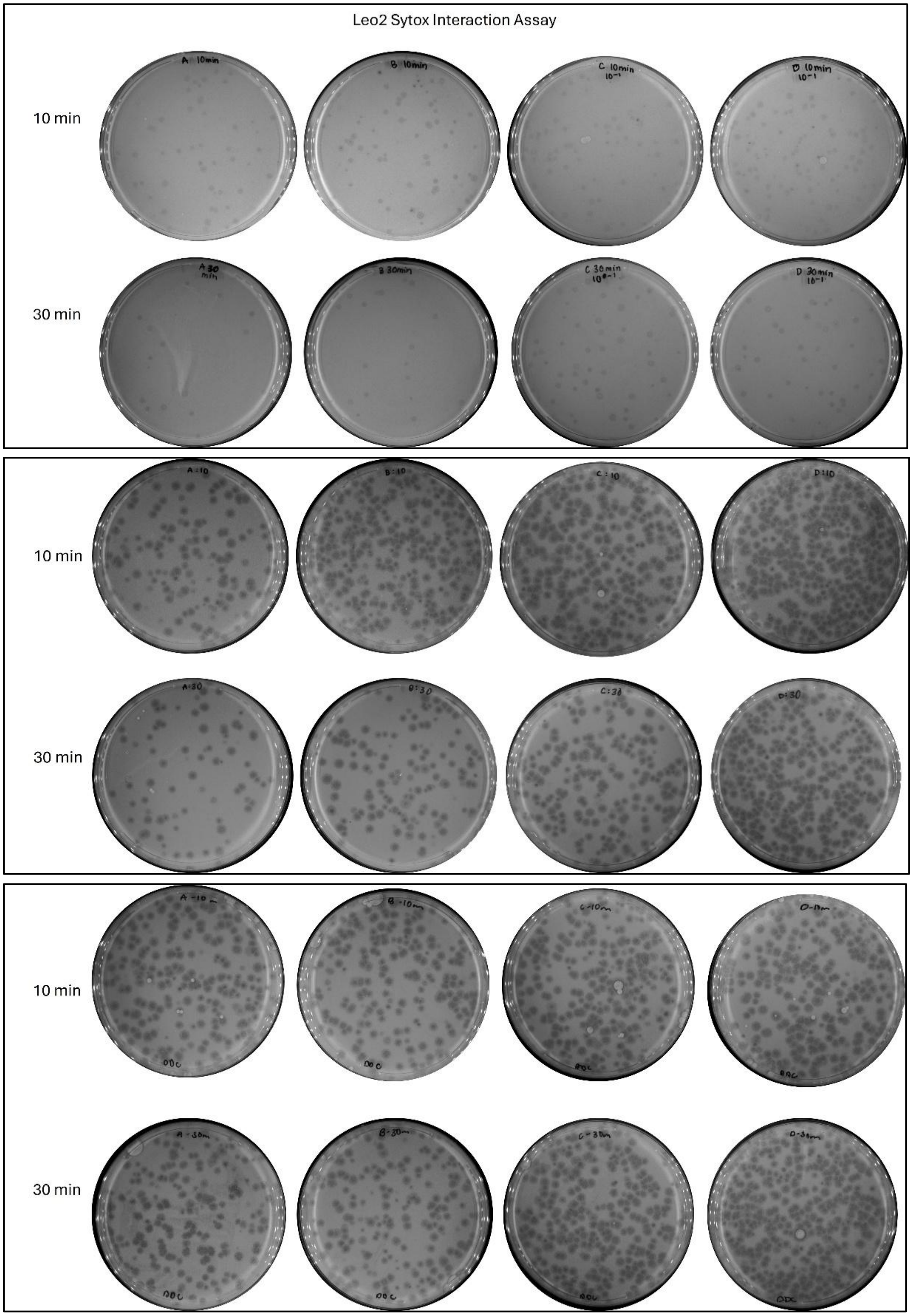

**Figure S5. Leo2 plaque formation following SYTOX exposure.**

Transmitted-light micrographs showing plaque formations following a Leo2 bacteriophage interaction with SYTOX for 10 min and 30 min across n=3 independent replicates.

T7

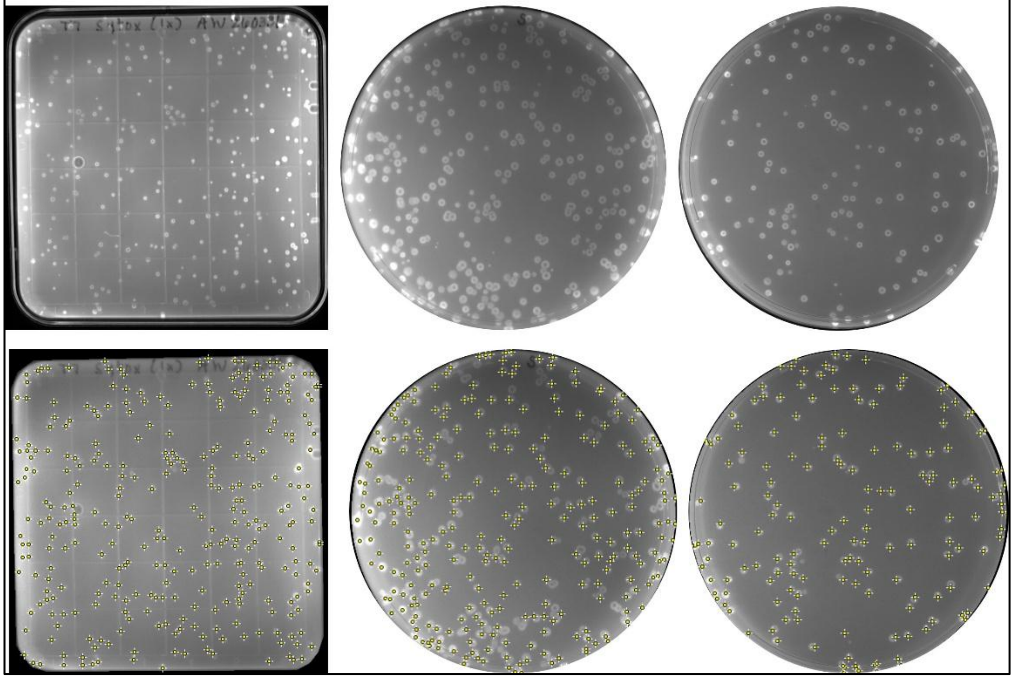

MS2

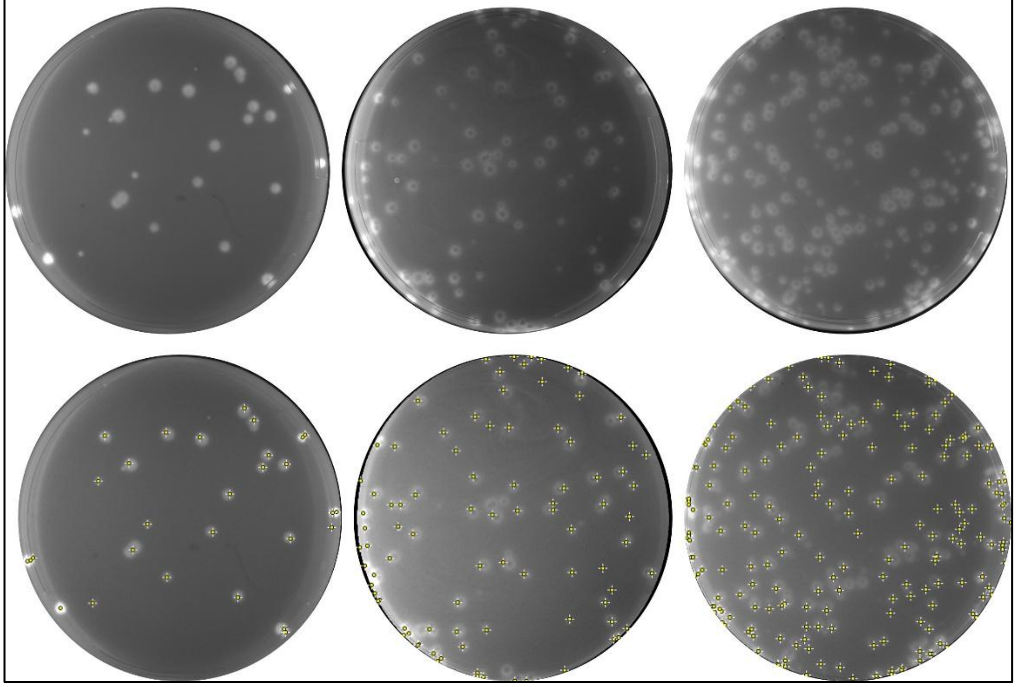

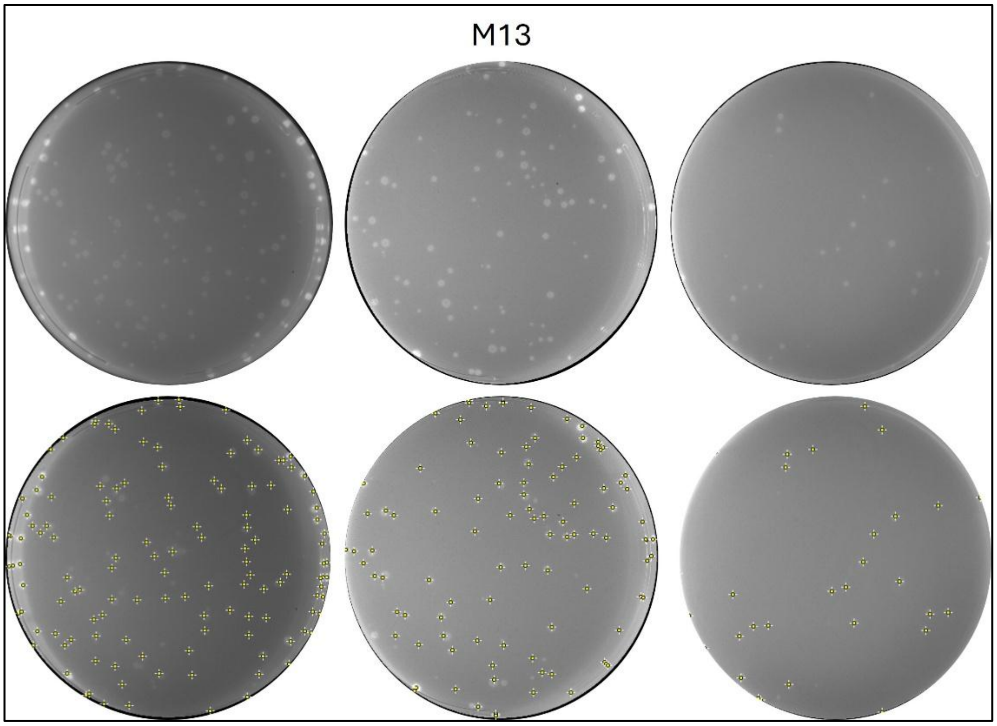

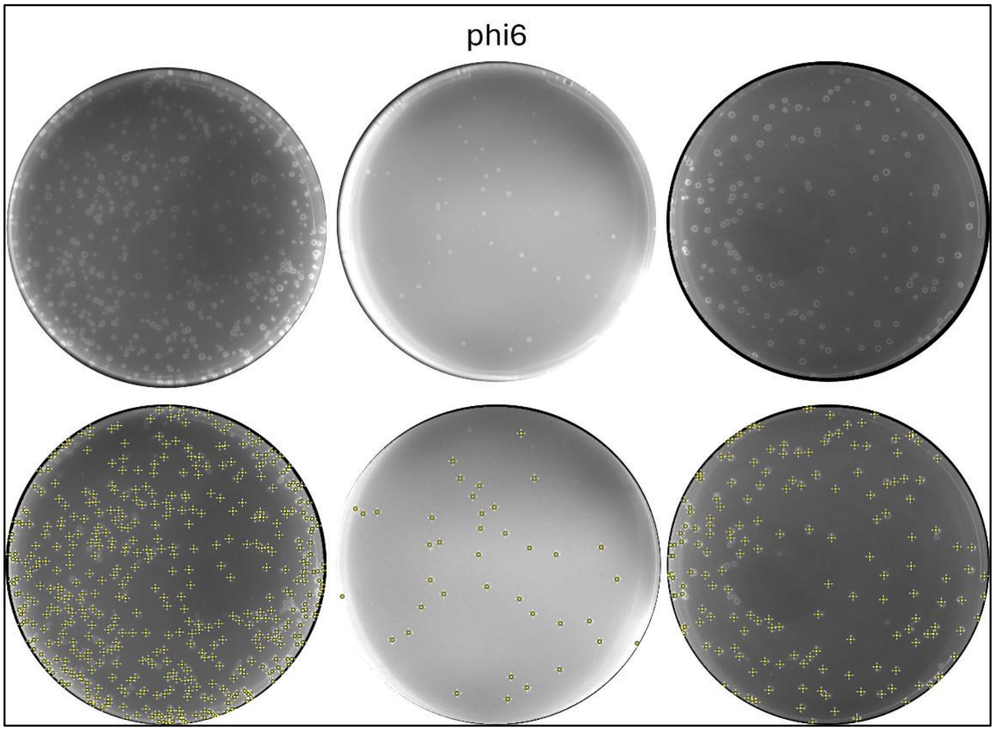

**Figure S6. Machine-based point selections across several bacteriophages.**

Fluorescent micrographs display accurate point selection with machine-based counting for T7, MS2, M13, and phi6 across n=3 independent replicates, as shown in Figure 2. Next day images were analyzed for MS2, M13, and phi6, whereas 6 h images were analyzed for T7.

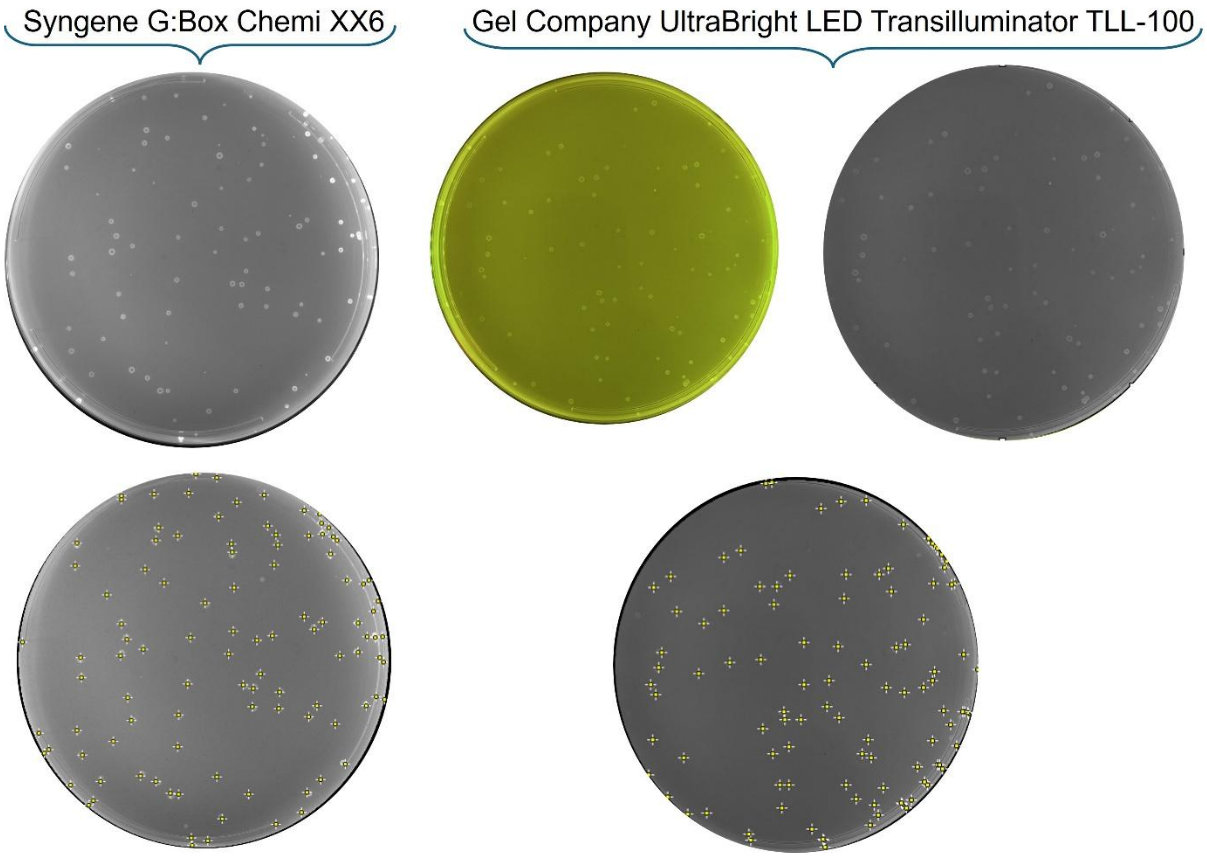

**Figure S7. SYTOX plaque imaging using an alternative blue-light imaging platform.**

Representative SYTOX-labeled plaques imaged on a Gel Company UltraBright LED Transilluminator TLL-100 using 470 nm blue LED illumination. Plaques remained sufficiently high-contrast for machine counting, indicating that SYTOX-based plaque detection is compatible with an alternative standard blue-light fluorescence imaging setup.

**Table S1. Manual and machine-based counts across bacteriophages over a time course**

Manual and machine-based counts were performed for SYTOX-labeled T7, MS2, M13, and phi6 plaques across a 24 h time course using three independent replicates. Conditions for which plates could not be counted are indicated with an "X". Conditions for which plates were not tested with machine-based counting are indicated with "N/T".

Image Time Course for T7, MS2, M13, and phi6 (3 replicates)

| Bacteriophage | Manual Count | Machine Count at 2 h | Machine Count at 4 h | Machine Count at 6 h | Machine Count at 8 h | Machine Count Next day (18-24 h) |
|---|---|---|---|---|---|---|
| T7 | 301 | X | 310 | 300 | 315 | 772 |
| T7 | 309 | X | 316 | 314 | 312 | 625 |
| T7 | 153 | X | 163 | 166 | 162 | 752 |
| MS2 | 26 | X | 42 | 30 | N/T | 29 |
| MS2 | 197 | X | 151 | 202 | N/T | 202 |
| MS2 | 83 | X | 90 | 91 | N/T | 91 |
| M13 | 92 | X | X | X | X | 100 |
| M13 | 125 | X | X | X | X | 133 |
| M13 | 26 | X | X | X | X | 30 |
| phi6 | 165 | X | X | X | X | 174 |
| phi6 | 34 | X | X | X | X | 38 |
| phi6 | 446 | X | X | X | X | 450 |

**Table S2. Machine-based counting across bacteriophage conditions**

Fluorescent micrographs for MS2, M13, and phi6 display accurate point selections with machine-based counting with next day images. T7 micrograph displays inaccurate counting at 24h and requires a fallback workflow to enable accurate machine-based counting.

| | T7 | | MS2 | | M13 | | phi6 | |
|---|---|---|---|---|---|---|---|---|
| | Manual | Machine | Manual | Machine | Manual | Machine | Manual | Machine |
| Replicate 1 | 301 | 300 | 26 | 29 | 125 | 133 | 446 | 450 |
| Replicate 2 | 274 | 284 | 197 | 202 | 92 | 100 | 165 | 174 |
| Replicate 3 | 153 | 166 | 83 | 91 | 26 | 30 | 34 | 38 |

**Table S3. Statistical analysis of the liquid-phase Leo2–*Bacillus safensis* SYTOX interaction assay.**

$Log_{10}$-transformed supernatant-recoverable Leo2 titers were analyzed by ordinary three-way ANOVA with factors of time, host-cell presence, and SYTOX presence, followed by Šídák's multiple-comparisons test for planned pairwise comparisons. Host-cell presence was the only significant source of variation, whereas SYTOX presence, time, and all interaction terms were not significant.

| Source of Variation | % of total variation | P value | P value summary | Significant? |
|---|---|---|---|---|
| Time (10-30 min) | 0.2271 | 0.5816 | ns | No |
| Host Presence | 87.17 | <0.0001 | **** | Yes |
| Sytox Presence | 0.008033 | 0.9171 | ns | No |
| Time (10-30 min) x Host Presence | 0.6479 | 0.3563 | ns | No |
| Time (10-30 min) x Sytox Presence | 0.1042 | 0.7082 | ns | No |
| Host Presence x Sytox Presence | 0.1226 | 0.6850 | ns | No |
| Time (10-30 min) x Host Presence x Sytox Presence | 0.2345 | 0.5756 | ns | No |
| **ANOVA table** | **SS** | **DF** | **MS** | **F (DFn, DFd)** |
| Time (10-30 min) | 0.07061 | 1 | 0.07061 | F (1, 16) = 0.3163 |
| Host Presence | 27.10 | 1 | 27.10 | F (1, 16) = 121.4 |
| Sytox Presence | 0.002497 | 1 | 0.002497 | F (1, 16) = 0.01119 |
| Time (10-30 min) x Host Presence | 0.2014 | 1 | 0.2014 | F (1, 16) = 0.9024 |
| Time (10-30 min) x Sytox Presence | 0.03241 | 1 | 0.03241 | F (1, 16) = 0.1452 |
| Host Presence x Sytox Presence | 0.03811 | 1 | 0.03811 | F (1, 16) = 0.1707 |
| Time (10-30 min) x Host Presence x Sytox Presence | 0.07290 | 1 | 0.07290 | F (1, 16) = 0.3266 |
| Residual | 3.571 | 16 | 0.2232 | |

| Šídák's multiple comparisons test | Mean diff. | 95.00% CI of diff. | Below threshold? | Summary | Adjusted P Value |
|---|---|---|---|---|---|
| Row 1:Leo2, Host, Sytox vs. Row 2:Leo2, Host, Sytox | 0.255 | -1.029 to 1.539 | No | ns | 0.9998 |
| Row 1:Leo2, Host vs. Row 2:Leo2, Host | 0.3284 | -0.9551 to 1.612 | No | ns | 0.9981 |
| Row 1:Leo2, LB Broth, Sytox vs. Row 2:Leo2, LB Broth, Sytox | 0.109 | -1.175 to 1.393 | No | ns | >0.9999 |
| Row 1:Leo2, LB Broth vs. Row 2:Leo2, LB Broth | -0.2585 | -1.542 to 1.025 | No | ns | 0.9998 |
| Row 1:Leo2, Host, Sytox vs. Row 1:Leo2, Host | -0.1368 | -1.420 to 1.147 | No | ns | >0.9999 |

| | | | | | |
|---|---|---|---|---|---|
| Row 1:Leo2, LB Broth, Sytox vs. Row 1:Leo2, LB Broth | 0.243 | -1.041 to 1.527 | No | ns | >0.9999 |
| Row 2:Leo2, Host, Sytox vs. Row 2:Leo2, Host | -0.06336 | -1.347 to 1.220 | No | ns | >0.9999 |
| Row 2:Leo2, LB Broth, Sytox vs. Row 2:Leo2, LB Broth | -0.1244 | -1.408 to 1.159 | No | ns | >0.9999 |
| Row 1:Leo2, Host, Sytox vs. Row 1:Leo2, LB Broth, Sytox | -2.132 | -3.416 to -0.8484 | Yes | *** | 0.0006 |
| Row 1:Leo2, Host vs. Row 1:Leo2, LB Broth | -1.752 | -3.036 to -0.4685 | Yes | ** | 0.004 |
| Row 2:Leo2, Host, Sytox vs. Row 2:Leo2, LB Broth, Sytox | -2.278 | -3.562 to -0.9944 | Yes | *** | 0.0003 |
| Row 2:Leo2, Host vs. Row 2:Leo2, LB Broth | -2.339 | -3.623 to -1.055 | Yes | *** | 0.0002 |
